# The peroxiredoxin Tsa1 extends the lifespan of budding yeast by maintaining the stability of the ribosomal RNA gene cluster

**DOI:** 10.1101/2024.03.14.585068

**Authors:** Junno Ohira, Mariko Sasaki, Takehiko Kobayashi

## Abstract

Tsa1 is a major budding yeast peroxiredoxin that also has nonperoxidase functions. Here we demonstrate that Tsa1 is required for stabilizing the ribosomal RNA gene (rDNA) cluster. Tsa1 deficiency led to lower replication initiation and an elevated recombination frequency in the rDNA. However, the absence of Tsa1 did not affect Fob1-dependent replication fork arrest at the replication fork barrier (RFB) site, indicating that forks at this site are arrested stably. Tsa1 deficiency did not affect the frequency of DNA double-strand breaks (DSBs) but enhanced transcription from the regulatory promoter E-pro toward the RFB. Because transcription from E-pro inhibits cohesin association and DSB end resection, elevated transcription may cause rDNA instability. We showed that the shortened lifespan and rDNA instability in the *tsa1*Δ mutant were largely suppressed by the *fob1* mutation. Therefore, Tsa1-mediated rDNA stabilization in response to replication fork arrest is important for prolonging lifespan. The *tsa1*Δ mutant accumulated single-strand breaks in the rDNA in a Fob1-independent manner. Taken together, these findings suggest that Tsa1 plays a crucial role in maintaining rDNA stability in both Fob1-dependent and Fob1-independent manners and contributes to lifespan extension.

## INTRODUCTION

Aging is the gradual deterioration of biological functions, which triggers various diseases, such as cancer, and ultimately leads to death (López-Otín et al., 2013; McHugh and Gil, 2018; Di Micco et al., 2021). Aging is influenced by various processes, including genome instability, dysregulation of nutrient sensing, mitochondrial dysfunction, and cellular senescence (López-Otín *et al*., 2013). The budding yeast *Saccharomyces cerevisiae* undergoes asymmetric cell division, in which a large mother cell produces a small daughter cell by budding (Mortimer and Johnston, 1959). Like human cells, budding yeast cells have a replicative lifespan and stop dividing after a finite number of times. Compared with that of higher eukaryotes, the replicative lifespan of budding yeast has a shorter division time, it can be determined by counting the number of daughter cells produced from mother cells, and genetic manipulation is easily performed. Therefore, budding yeast has been used as a model organism for studying the molecular mechanisms underlying cellular senescence.

The ribosomal RNA gene (rDNA) is a highly abundant repetitive sequence in eukaryotic genomes. Budding yeast normally has ∼150 rDNA copies, which are tandemly repeated at a single locus on chr XII. In addition to 35S and 5S rDNA, each copy contains an origin of DNA replication, a replication fork barrier (RFB) site, and a bidirectional, RNA polymerase II promoter, E-pro. After DNA replication is initiated, the replication fork that moves in a direction opposite to that of 35S rDNA is arrested at the RFB site by the Fob1 protein, leading to DNA double-strand breaks (DSBs) (Fig. 1A) (Brewer et al., 1992; Kobayashi et al., 1992; Kobayashi and Horiuchi, 1996; Burkhalter and Sogo, 2004; Kobayashi et al., 2004). End resection of these DSBs, an initiating event for homologous recombination (HR), is normally restricted. We previously demonstrated that removal of this suppression, for example, by the absence of the histone deacetylase Sir2 and by transcription from E-pro, induces HR-mediated DSB repair (Fig. 1A) (Sasaki and Kobayashi, 2017; Sasaki and Kobayashi, 2023b; Sasaki and Kobayashi, 2023a). The use of the rDNA copy at an aligned position on the sister chromatid during DSB repair leads to the maintenance of rDNA copy number. However, when the copy at a misaligned position on the sister chromatid or another copy on the same chromosome is used, DSB repair can lead to rDNA instability accompanied by amplification or loss of rDNA copies. Cohesin complexes that are recruited to intergenic regions promote the use of aligned rDNA copies during DSB repair, but their binding is inhibited by transcription from E-pro (Fig. 1A) (Kobayashi *et al*., 2004; Kobayashi and Ganley, 2005; Sasaki and Kobayashi, 2023b). The histone deacetylase Sir2 plays an important role in the maintenance of rDNA stability by repressing transcription from E-pro to facilitate stable cohesin binding (Kobayashi *et al*., 2004; Kobayashi and Ganley, 2005).

**Figure 1.**
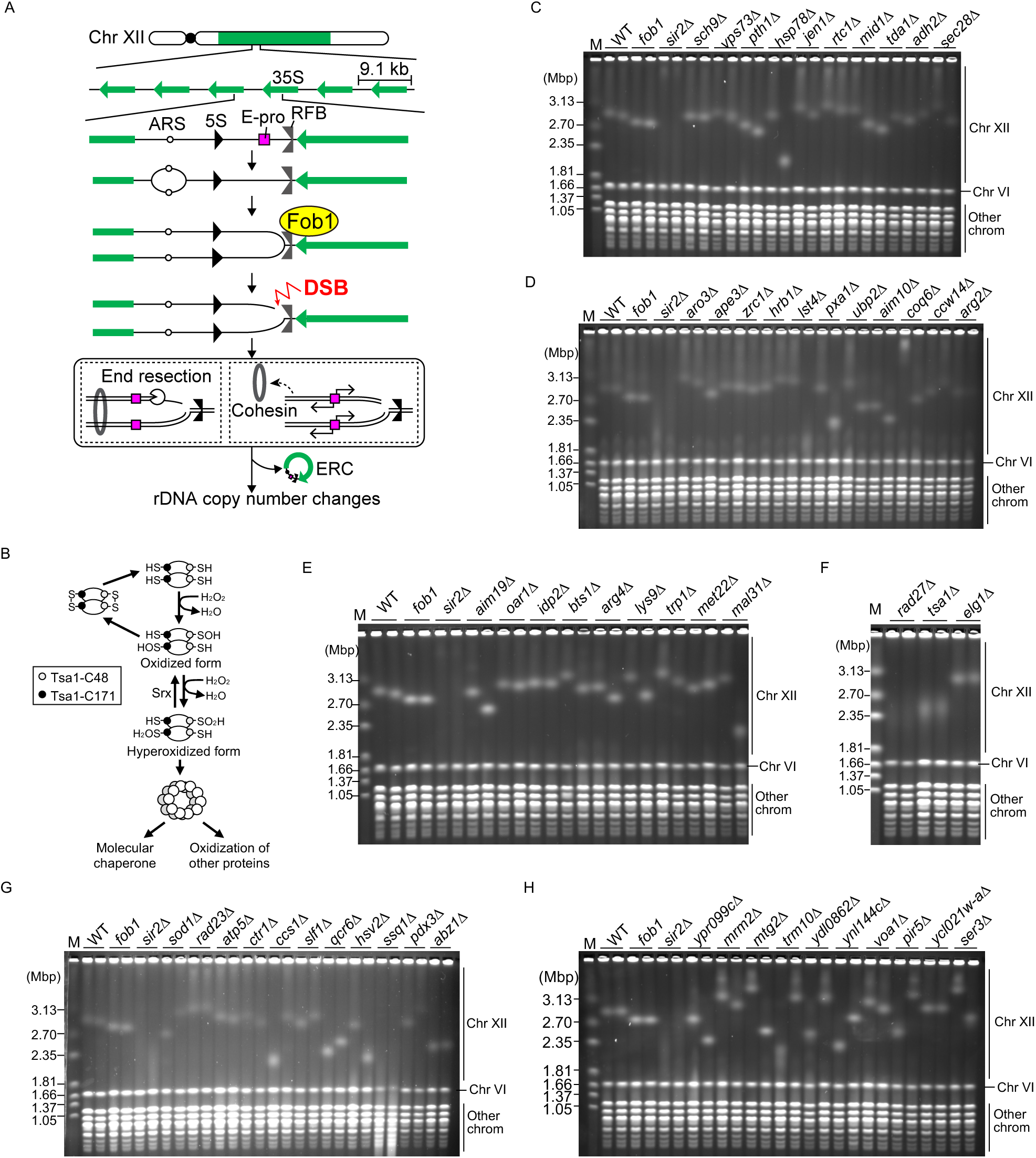
PFGE analysis of rDNA instability in the candidate strains. **(A)** rDNA copy number changes in budding yeast. 35S and 5S, (pre)rRNA transcription units; ARS, an origin of DNA replication; E-pro, an RNA polymerase II-dependent, bidirectional promoter that synthesizes noncoding RNA; RFB, replication fork barrier; DSB, DNA double-strand break; ERC, extrachromosomal rDNA circle. **(B)** The catalytic cycle of Tsa1. The ellipses are Tsa1 proteins. **(C–H)** Genomic DNA was prepared from two independent clones of the indicated mutant strains and separated by PFGE. Genomic DNA was also prepared from the WT, *fob1* and *sir2*Δ mutants and analyzed in parallel. DNA was stained with ethidium bromide. M indicates *H. wingei* chromosomal DNA markers.

The stability of the rDNA region has a substantial impact on the cellular senescence of budding yeast. When a *FOB1* gene is mutated, it not only stabilizes the rDNA but also extends the replicative lifespan (Defossez et al., 1999; Takeuchi et al., 2003; Menzel et al., 2014). The absence of Sir2 causes rDNA instability and shortens the replicative lifespan, while the overexpression of *SIR2* results in an extended lifespan (Sinclair and Guarente, 1997; Sinclair et al., 1997; Kaeberlein et al., 1999; Kobayashi *et al*., 2004; Saka et al., 2013). Therefore, the maintenance of rDNA stability is important for delaying cellular senescence.

Reactive oxygen species (ROS) are highly reactive molecules that are derived from oxygen and are composed of superoxide anion radicals, hydrogen peroxide (H_2_O_2_), and hydroxyl radicals [reviewed by (Sies and Jones, 2020)]. When present in excess, ROS are harmful because they can damage DNA, proteins and lipids. Damaged DNA induces a wide spectrum of mutations and chromosomal rearrangements (Huang et al., 2003). Some ROS act as signaling molecules and regulate various cellular processes, such as proliferation and differentiation, inflammation, tumorigenesis, cellular senescence and cell death, according to cellular ROS levels [reviewed in (Sies and Jones, 2020)]. Peroxiredoxins (Prxs) are conserved enzymes that convert H_2_O_2_ into H_2_O [reviewed in (Wood et al., 2003; Nystrom et al., 2012; Perkins et al., 2015)]. Tsa1 is a major Prx in budding yeast that acts as an obligate homodimer and preferentially forms a decamer to scavenge H_2_O_2_ [reviewed in (Perkins *et al*., 2015)]. Tsa1 oxidizes the thiol group (-SH) of cysteine (Cys) at position 48 to sulfenic acid (-SOH) while reducing H_2_O_2_ to H_2_O (Fig. 1B). The oxidized Tsa1 forms a disulfide bond with Cys171 of its partner Tsa1. Alternatively, oxidized Tsa1 further oxidizes sulfenic acid to the sulfinic acid (-SOOH) of Cys48 (the hyperoxidized form) while reducing H_2_O_2_ to H_2_O (Fig. 1B) (Wood *et al*., 2003). Hyperoxidized Tsa1 is inactive for peroxidase activity and needs to be reduced by sulfiredoxin Srx in budding yeast (Perkins *et al*., 2015). ROS induce damage to proteins, inducing their misfolding and aggregation. Hyperoxidized Tsa1 forms a double decamer or even a higher molecular weight form, which switches the function of Tsa1 from that of peroxidase to that of a molecular chaperone that mediates the disaggregation of proteins by recruiting Hsp70 and Hsp104 to misfolded proteins (Jang et al., 2004; Hanzen et al., 2016). Tsa1 is also involved in redox-dependent signal transduction through the oxidation of other proteins to regulate various cellular processes, such as gluconeogenesis (Delaunay et al., 2002; Sobotta et al., 2015; Irokawa et al., 2016).

Tsa1 deficiency not only leads to an increased mutation rate but also shortens the replicative lifespan (Huang *et al*., 2003; Molin et al., 2011). An increase in the dosage of Tsa1 extends the replicative lifespan but does not affect the mutation rate (Hanzen *et al*., 2016). Thus, the ability of Tsa1’s Prx activity to prevent mutations is less likely to be important for lifespan extension. However, previous studies have suggested that Tsa1 delays cellular aging through its chaperone function to facilitate the clearance of H_2_O_2_-mediated protein aggregates and its ability to modify protein kinase A in the nutrient signaling pathway (Hanzen *et al*., 2016; Roger et al., 2020).

In this study, we screened for factors involved in ROS production, the maintenance of rDNA stability and replicative lifespan and identified Tsa1 as a candidate factor. We found that cells lacking Tsa1 showed destabilized rDNA in a manner mostly dependent on Fob1. The *tsa1*Δ mutant showed lower efficiency of firing of the DNA replication origin in the rDNA, but the level of arrested forks was comparable between the WT and *tsa1*Δ mutant, demonstrating that forks stalled at the RFB were stably arrested. The *tsa1*Δ mutant did not exhibit an altered level of DSBs. Tsa1 deficiency led to a greater level of transcripts synthesized from E-pro toward the RFB site, indicating that DSBs at the RFB site are prone to induce rDNA copy number changes, possibly due to the inhibition of cohesin binding and the induction of DSB end resection. In the *tsa1*Δ mutant, single-strand breaks (SSBs) accumulated near the 5S rDNA, regardless of the presence or absence of Fob1, which may also be responsible for rDNA instability. Last, previous studies have demonstrated that defects in proteostasis in the *tsa1*Δ mutant cause a shortened lifespan. Our finding that rDNA instability and lifespan shortening were suppressed by the deletion of *TSA1* suggests that Tsa1-mediated rDNA stabilization also contributes to lifespan extension. Because rDNA stabilization depends on Cys48 and Cys171, which are important for peroxidase function and redox-dependent target modulation, we propose that Tsa1 stabilizes rDNA by activating proteins involved in rDNA maintenance.

## RESULTS

### Tsa1 is required for the maintenance of the stability of the rDNA region

We aimed to identify factors that regulate rDNA stability, ROS levels, and replicative lifespan. Because ROS are primarily produced in the mitochondrial electron transport chain, mitochondrial dysfunction escalates ROS (Fang and Beattie, 2003; Seo et al., 2010). Aging yeast cells experience mitochondrial dysfunctions, but these defects can be prevented by suppressing vacuolar acidity and abolishing neutral amino acid transport (Hughes and Gottschling, 2012). We previously identified 713 gene knockout mutants that exhibit rDNA instability (Saka et al., 2016). Among these genes, we searched for those whose mutations are associated with altered lifespan, mitochondrial genome maintenance abnormalities, alterations in pH resistance, changes in oxidative stress resistance, and abnormal mitochondrial morphology and for genes associated with the Gene Ontology terms, cellular amino acid metabolic process, vesicle organization, mitochondria and vesicles, leading to the identification of 171 genes. Among them, 103 genes had already been characterized in previous studies or found in our preliminary analyses that their mutants did not show reproducibility in rDNA instability (Kobayashi and Sasaki, 2017; Hosoyamada et al., 2019; Goto et al., 2021). Thus, we aimed to construct mutants lacking each of the remaining 68 genes and analyze their rDNA instability. Fourteen genes could not be deleted in the W303 strain background for unknown reasons. Thus, the involvement of the remaining 54 genes in the maintenance of rDNA stability was examined.

To assess rDNA stability, we isolated genomic DNA from two independent clones of candidate mutants and separated the DNA by pulsed field gel electrophoresis (PFGE). We then examined the size heterogeneity of chr XII, which carries the rDNA array. We included the WT strain, the *fob1* mutant that displayed a sharper chr XII band due to the suppression of rDNA copy number changes, and the *sir2*Δ mutant that displayed a smeared chr XII band due to continuous rDNA copy number changes. Two clones of the *sch9*Δ mutant displayed chr XII bands that were as sharp as those observed in the WT clones (Fig. 1C), demonstrating that the rDNA region was stable in this mutant. Most of the mutants analyzed exhibited stable rDNA. Two clones of the *hsp78*Δ mutant showed different sizes of chr XII: one clone had chr XII of ∼ 2.7 Mbp, and the other clone had chr XII of <2.35 Mbp (Fig. 1C). Although their sizes differed among the two clones, both displayed chr XII bands that were as sharp as those in the WT. Yeast cells can undergo spontaneous rDNA copy number changes during transformation procedures that are performed during strain construction (Kwan et al., 2016). The *hsp78*Δ mutant most likely experienced such rDNA copy number changes during strain construction but was capable of stably maintaining that copy number. Mutants such as *pxa1*Δ (Fig. 1D), *coq6*Δ (Fig. 1D), *mal31*Δ (Fig. 1E), *ccs1*Δ (Fig. 1G), and *mtg2*Δ (Fig. 1H) exhibited similar patterns. We concluded that genes deleted in these mutants were not involved in the maintenance of rDNA stability. In the *ssq*Δ mutant, chr XII bands were barely observed as distinct bands (Fig. 1G). We attempted to confirm this phenotype by generating a diploid strain heterozygous for *ssq1*Δ, isolating haploid *ssq1*Δ clones by tetrad analysis and analyzing their rDNA stability. We failed to obtain viable haploid *ssq1*Δ spores in the W303 strain background. Thus, this gene was not analyzed further.

The *rad27*Δ and *tsa1*Δ mutants exhibited chr XII bands that were smeared compared to those of the WT and as smeared as those of the *sir2*Δ mutant (Fig. 1F). *RAD27* encodes a structure-specific nuclease that plays important roles in maintaining the integrity of nuclear genomes. Rad27 is also localized in the mitochondria and is involved in maintaining mitochondrial DNA stability (Kaniak-Golik et al., 2017). *TSA1* encodes a Prx that scavenges H_2_O_2_, which suppresses the accumulation of mutations (Huang *et al*., 2003). Tsa1 also performs nonperoxidase functions, such as maintaining proteostasis (Hanzen *et al*., 2016; Irokawa *et al*., 2016). The absence of *TSA1* causes a shortened replicative lifespan, while the overexpression of *TSA1* extends the lifespan (Molin *et al*., 2011; Hanzen *et al*., 2016). Because Tsa1 is more directly involved in controlling H_2_O_2_ and regulates cellular aging, we sought to further examine how Tsa1 regulates rDNA stability and replicative lifespan.

### Tsa1, but not other ROS regulators, is important for promoting rDNA stability

Budding yeast has five Prxs, Tsa1, Tsa2, Ahp1, Prx1, and Dot5. Tsa1, Tsa2 and Ahp1 are localized in the cytoplasm, while Prx1 is localized in mitochondria, and Dot5 is localized in the nucleus (Park et al., 2000). Tsa1 is a major cytosolic Prx that constitutes approximately 90% of the total intracellular Prx (Tairum et al., 2016). The *tsa1*Δ mutant showed a smeared chr XII band, but mutants lacking other Prxs showed stable rDNA, as their chr XII bands were as sharp as those in WT cells (Fig. 2A, 2B).

**Figure 2.**
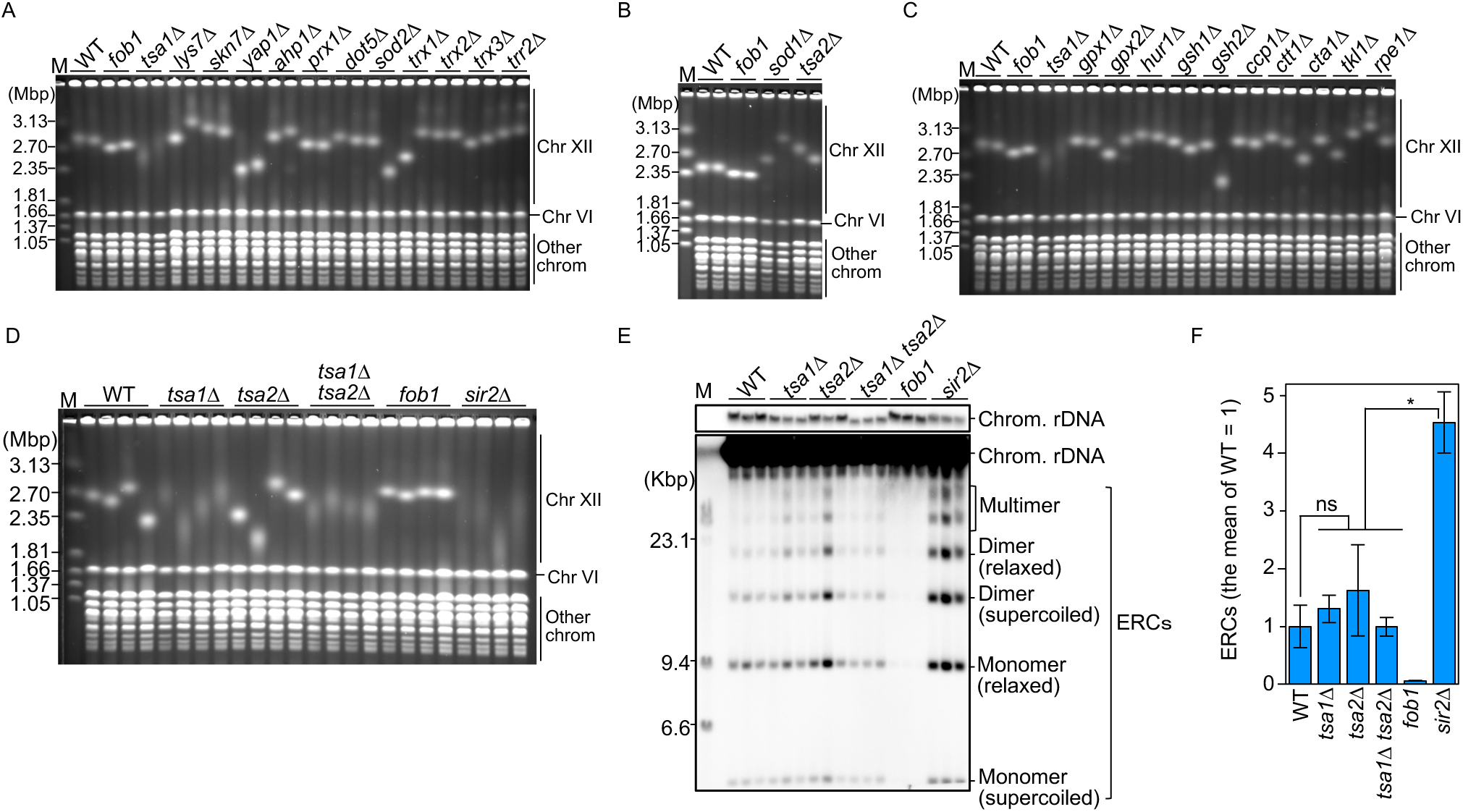
Tsa1 is important for the maintenance of rDNA stability. **(A–D)** PFGE analysis of the size heterogeneity of chr XII. The indicated genes that encode enzymes that remove ROS were deleted from WT cells. DNA was extracted from two independent clones of the WT and all mutant strains and separated by PFGE. DNA was stained with ethidium bromide. M indicates *H. wingei* chromosomal DNA markers. **(E)** ERC detection. DNA was isolated from two independent clones of the indicated strains and separated by agarose gel electrophoresis, followed by Southern blotting with rDNA probe 1, as indicated in Fig. 4A. The chromosomal rDNA array and the supercoiled and relaxed forms of monomeric and dimeric ERCs are indicated. The sizes of lambda DNA-HindIII markers are indicated. **(F)** Quantitation of ERCs. The ERCs in (E) were quantified. The bars show the means ± s.e.m. Multiple comparisons were performed by one-way ANOVA, followed by Tukey’s multiple comparisons test; * indicates a statistically significant difference (p ≤ 0.05); ns indicates no significant difference.

Budding yeast possesses other enzymes that function to remove ROS (Huang *et al*., 2003; Thorpe et al., 2004; Perrone et al., 2008). For instance, Sod1 and Sod2 function as superoxide dismutases that convert superoxide anion radical O_2_^−^ into H_2_O_2_ and O_2_. Catalases such as Ctt1 and Cta1 decompose H_2_O_2_ into H_2_ and dH_2_O, while the glutathione peroxidases Gpx1, Gpx2 and Hyr1 reduce H_2_O_2_ to H_2_O and O_2_ in the presence of glutathione. Additionally, the thioredoxins Trx1, Trx2, Trx3 and Trr2 reduce disulfide bonds formed by the cysteine residues of peroxidases. Other oxidant defense genes, such as the transcription factors Yap1 and Skn7, are also part of the oxidant response system. The absence of any of these enzymes did not destabilize rDNA (Fig. 2A–2C).

Tsa1 and Tsa2 are highly homologous. The *tsa2*Δ single mutant does not show the mutator phenotype, but the *tsa1*Δ *tsa2*Δ double mutant not only displays higher mutation rates but also becomes sensitive to hydrogen peroxide compared to the *tsa1*Δ single mutant (Huang *et al*., 2003; Wong et al., 2004; Iraqui et al., 2009). Moreover, the *TSA2* expression level is increased in the *tsa1*Δ mutant (Wong et al., 2002). These findings indicate that Tsa2 can confer resistance to hydrogen peroxide and prevent the accumulation of mutations when Tsa1 is absent. To determine whether Tsa2 functions to promote rDNA stability in the *tsa1*Δ mutant, we generated a diploid strain that was heterozygous for *tsa1*Δ and *tsa2*Δ, isolated haploid clones by tetrad dissection and analyzed the rDNA stability of four independent clones—the WT, *tsa1*Δ, *tsa2*Δ, and *tsa1*Δ tsa2Δ— by PFGE. Three out of the four *tsa2*Δ clones exhibited chr XII bands that were as sharp as those in the WT clones (Fig. 2D). The second *tsa2*Δ clone had a smaller chr XII, and its band was smeared. When the rDNA copy number is substantially reduced from the normal copy number of ∼150, cells enter a phase to induce rDNA amplification (Kobayashi et al., 1998). We speculate that the second *tsa1*Δ clone inherited chr XII with a substantially low rDNA copy number during sporulation and was in the rDNA amplification phase. Based on the phenotypes of the other three *tsa2*Δ clones, we concluded that the *tsa2*Δ mutation did not compromise rDNA stability. The *tsa1*Δ *tsa2*Δ mutant clones were as smeared as those of the *tsa1*Δ single mutant (Fig. 2D).

Chromosomal rDNA copy number changes are often associated with the production of ERCs, which can be used as an indicator of rDNA instability (Fig. 1A) (Kaeberlein *et al*., 1999; Kobayashi *et al*., 2004). Compared with WT cells, the *sir2*Δ mutant showed an increase in the level of ERCs (Fig. 2E, 2F), consistent with previous findings (Kaeberlein *et al*., 1999; Hosoyamada *et al*., 2019; Goto *et al*., 2021; Yokoyama et al., 2023). Although the *tsa1*Δ mutant displayed chromosomal rDNA instability to a degree comparable to that of the *sir2*Δ mutant, its level of ERCs was comparable to that in WT cells. Compared with those in WT cells, the ERC levels in the *tsa2*Δ and *tsa1*Δ *tsa2*Δ double mutants did not increase (Fig. 2F). Therefore, both PFGE and ERC analyses demonstrated that deletion of *TSA2* did not exacerbate rDNA instability in the *tsa1*Δ single mutants. These findings suggest that among the various factors involved in the metabolism of ROS, Tsa1 plays a uniquely important role in promoting rDNA stability.

### Tsa1 suppresses rDNA instability in response to Fob1-mediated replication fork arrest

To understand how Tsa1 promotes rDNA stability, we examined whether Tsa1 functions in a major rDNA-destabilizing pathway that depends on Fob1-mediated replication fork arrest at the RFB. To this end, we constructed a diploid strain heterozygous for *tsa1*Δ and *fob1*, isolated haploid clones by tetrad dissection, and analyzed rDNA stability in four independent clones of WT, *fob1*, *tsa1*Δ, and *tsa1*Δ *fob1* by PFGE. Compared with the other three clones, the WT clones showed stable rDNA, except for the fourth clone, which had a smaller and smeared chr XII band (Fig. 3A). We speculate that this clone had spontaneously lost rDNA copies and entered the rDNA amplification phase. The chr XII bands of four *fob1* clones were sharper than those of the WT clones (Fig. 3A). The *tsa1*Δ mutant showed smeared chr XII bands, but this smearing was largely suppressed by the introduction of a *fob1* mutation. However, the chr XII bands were still more smeared than those in the *fob1* single mutant (Fig. 3A). This phenotype was in contrast to that of the *sir2*Δ mutant, in which smearing of the chr XII bands was nearly completely suppressed by the introduction of a *fob1* mutation to the degree seen in the *fob1* single mutant.

**Figure 3.**
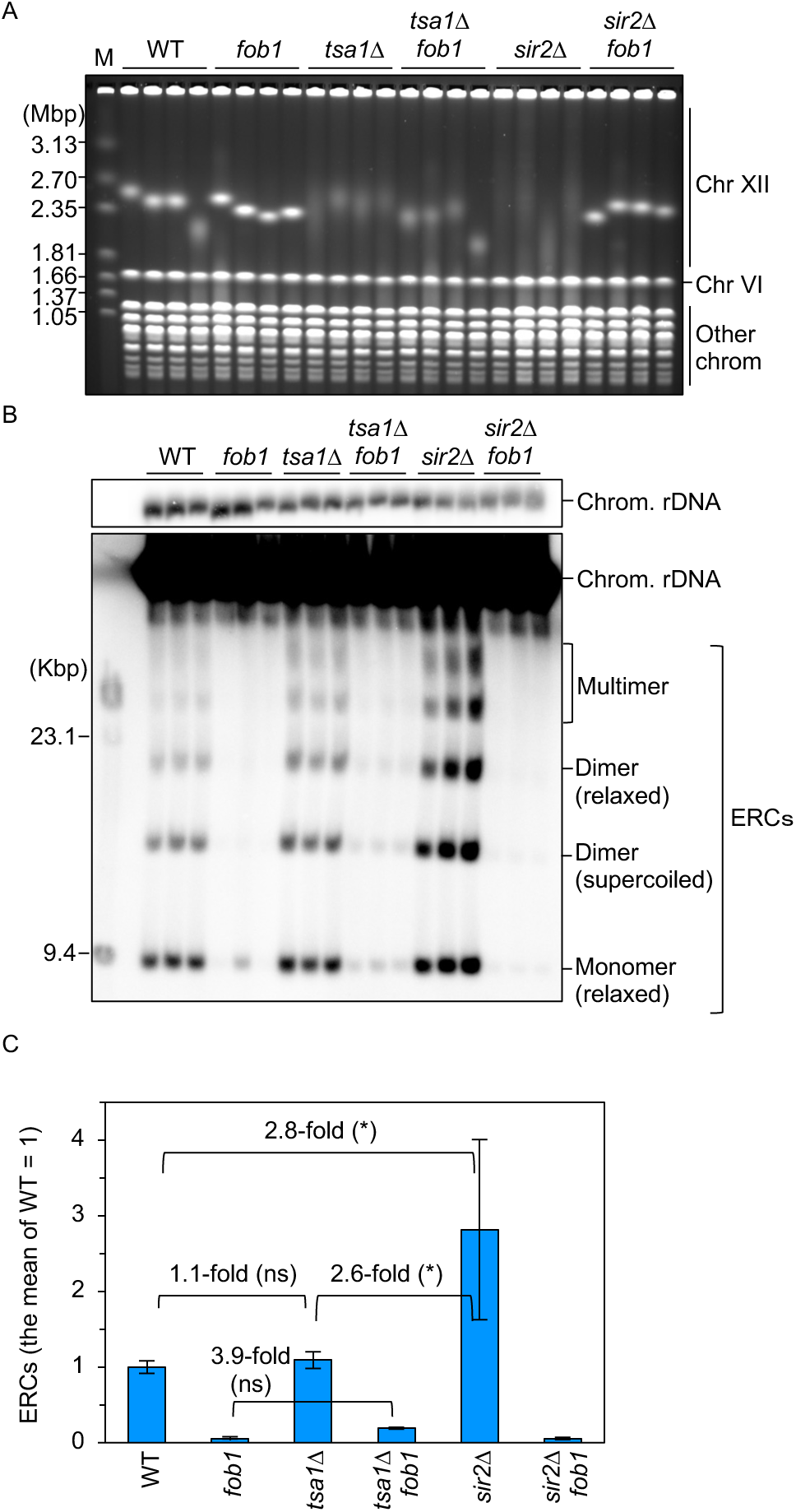
Tsa1 mainly acts in a Fob1-dependent pathway to stabilize rDNA. **(A)** PFGE analysis of the size heterogeneity of chr XII. DNA was extracted from four independent clones of the indicated strains and separated by PFGE. DNA was stained with ethidium bromide. M indicates *H. wingei* chromosomal DNA markers. **(B)** ERC detection. DNA was isolated from three independent clones of the indicated strains and separated by agarose gel electrophoresis, followed by Southern blotting with rDNA probe 1, as shown in Fig. 4A. The chromosomal rDNA array and the supercoiled and relaxed forms of monomeric and dimeric ERCs are indicated. The sizes of lambda DNA-HindIII markers are indicated. **(C)** Quantitation of ERCs. The ERCs in (B) were quantified. The bars show the means ± s.e.m. Multiple comparisons were performed by one-way ANOVA, followed by Tukey’s multiple comparisons test; * indicates a statistically significant difference (p ≤ 0.05); ns indicates no significant difference.

We also assessed rDNA stability by ERC analysis. The level of ERCs in the *tsa1*Δ mutant was comparable to that in the WT (Fig. 3B, 3C). The absence of Fob1 in the *sir2*Δ mutant background suppressed ERC production to a level comparable to that in the *fob1* mutant. The level of ERCs in the *tsa1*Δ *fob1* mutant was ∼3.9-fold greater than that in the *fob1* mutant, although this difference was not statistically significant according to multiple comparisons among all the strains (Fig. 3B, 3C). Taken together, these findings suggest that the function of Tsa1 is mainly to stabilize chromosomal rDNA copy numbers in response to Fob1-dependent replication fork arrest, but Tsa1 also contributes to the maintenance of rDNA stability through its Fob1-independent functions.

### Tsa1 promotes efficient replication initiation and suppresses recombination in rDNA

To understand how Tsa1 regulates the cellular response to Fob1-dependent replication fork arrest, we examined the pattern of DNA replication and the frequency of recombination in the rDNA region in the *tsa1*Δ mutant. We isolated genomic DNA from exponentially growing WT, *tsa1*Δ, *fob1*, and *tsa1*Δ *fob1* cells, digested it with the restriction enzyme NheI, which has two recognition sites within each rDNA copy, and performed two-dimensional (2D) agarose gel electrophoresis (Fig. 4A). DNA was first separated by agarose gel electrophoresis according to molecular mass in the first dimension and then according to molecular mass and shape in the second dimension, followed by Southern blotting with rDNA probe 1 (Fig. 4A, 4B).

**Figure 4.**
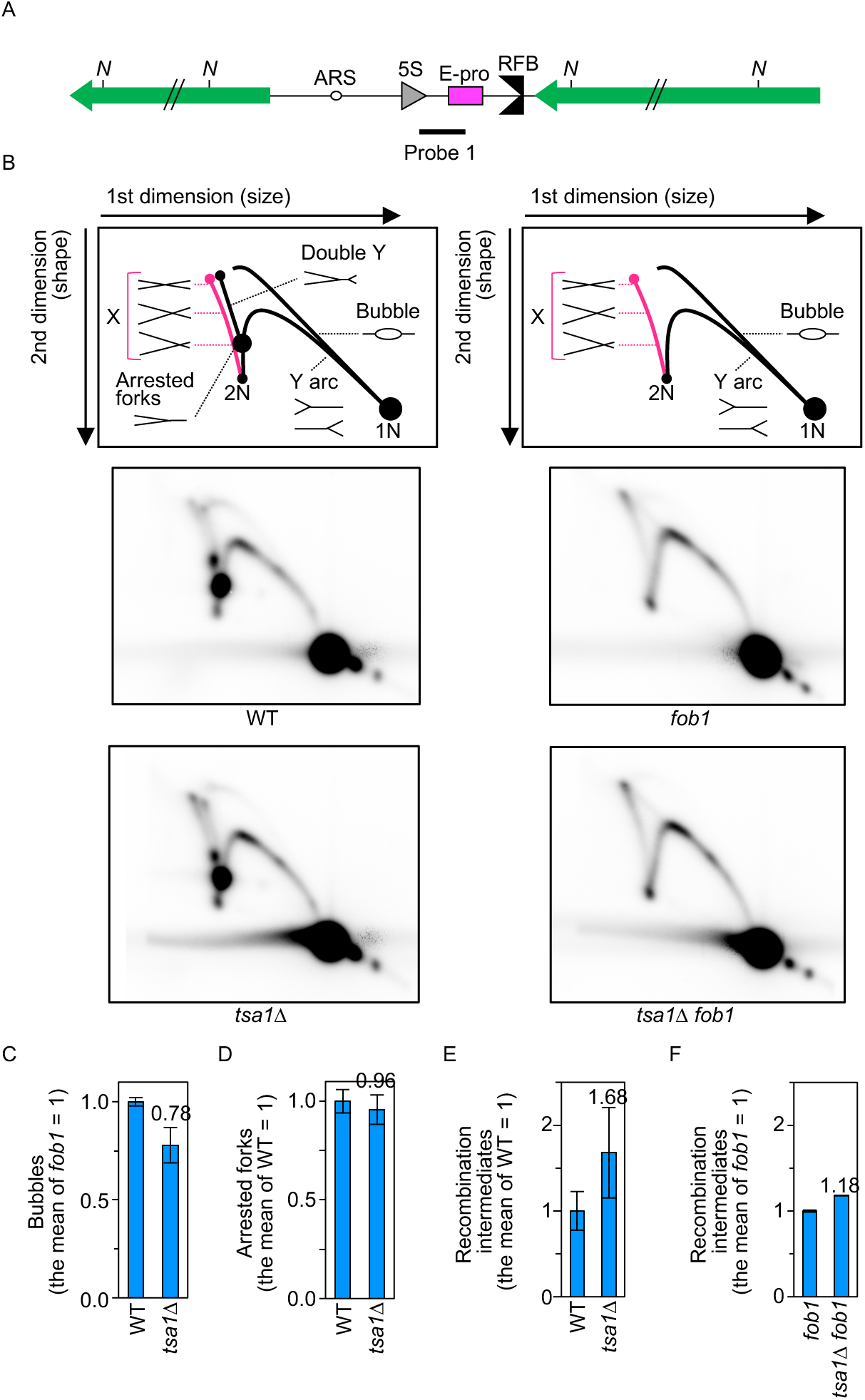
Tsa1 suppresses recombination mainly in a Fob1-dependent manner. **(A)** Restriction map of the probe used for 2D analysis. The restriction sites for NheI (N) and the position of probe 1 are indicated. **(B)** 2D agarose gel electrophoresis. Genomic DNA from the indicated strains was digested with NheI and separated according to size in the first dimension and according to size and shape in the second dimension, followed by Southern blotting with the probe, as indicated in (A). The diagrams on the left and right show the expected migration patterns of different replications of the *FOB1* and *fob1* strains, respectively. 1N represents linear DNA, and 2N represents linear DNA that was nearly fully replicated. X structures that include recombination intermediates are indicated in pink. **(C–F)** Quantitation of DNA replication and recombination intermediates. The frequencies of bubbles, arcs (C), arrested forks (D), X structures in the *FOB1* background (E) and X structures in the *fob1* background (F) were determined by quantifying the signal of each intermediate relative to the total number of replication intermediates. The levels of different molecules in each mutant were normalized to the average of the WT clones (C-E) and *fob1* clones (F). Bars indicate the range of two independent experiments.

DNA molecules where DNA replication is initiated from a replication origin (ARS) are bubble-shaped molecules, while DNA molecules where DNA replication progresses passively through the NheI fragment generate a Y arc (Fig. 4B). DNA replication forks that move rightward from a replication origin produce an intense signal along the Y arc. DNA molecules where the other fork approaches an arrested fork at the RFB from a downstream origin are detected as double-Y molecules, which protrude from the spot of arrested forks at the RFB (Fig. 4B, left). The frequency of origin firing was assessed by determining the proportion of the signal for bubble-shaped molecules relative to the total signal for replication intermediates. The replication initiation frequency in the *tsa1*Δ mutant decreased by ∼20% relative to that in the WT (Fig. 4C). More than 90% of replication forks that progress from a fired replication origin are stalled at the RFB site (Brewer *et al*., 1992). Considering that fewer origins fired in the *tsa1*Δ mutant, fewer forks would be arrested at the RFB site in this mutant than in WT cells. Unexpectedly, the number of arrested forks was comparable between the WT and *tsa1*Δ strains (Fig. 4D). Thus, arrested forks stay longer, most likely because it takes longer for the converging forks to arrive and release arrested forks.

X-shaped molecules that protrude from the 2N spot include recombination intermediates, such as Holliday unction structures (Fig. 4B). The levels of these molecules, which were estimated by the proportion of their signal relative to the total number of replication intermediates, were ∼1.7-fold greater than those in WT cells (Fig. 4E). In the *fob1* and *tsa1*Δ *fob1* mutants, replication forks were not arrested at the RFB, and double-Y molecules were not formed in these mutants (Fig. 4B, right). X-shaped molecules were only slightly increased by the deletion of *TSA1* in the *fob1* background (Fig. 4F). Taken together, these findings indicate that Tsa1 suppresses recombination primarily in response to Fob1-mediated replication fork arrest.

### Tsa1 negatively regulates the transcription of noncoding RNA from E-pro toward RFB

Arrested forks at the RFB are broken into DSBs (Weitao et al., 2003; Burkhalter and Sogo, 2004; Kobayashi *et al*., 2004). End resection of DSBs initiates homologous recombination, which is prone to induce rDNA copy number changes (Sasaki and Kobayashi, 2017; Sasaki and Kobayashi, 2023b; Sasaki and Kobayashi, 2023a). To analyze the frequency and repair of DSBs, we isolated genomic DNA from exponentially growing cells, digested DNA with the restriction enzyme Bgl II, and separated DNA by conventional agarose gel electrophoresis, followed by Southern blotting with indirect end-labeling (Fig. 5A). DNA molecules above the 8.5 kbp DNA size marker corresponded to large Y molecules that appeared descending from the apex of the Y arc toward the 2N signal in the 2D analyses (Fig. 5B). When Fob1 was present, it induced a strong fork arrest signal on the Y arc, producing a much stronger signal than that in the *fob1* mutant (Fig. 5B). When arrested forks at the RFB were broken, one-ended DSBs of ∼2.3 kbp in length were generated (Fig. 5A, 5B). The frequency of DSBs, which was determined by calculating the ratio of DSBs relative to Y-arc molecules, including arrested forks, was comparable between the WT and *tsa1*Δ mutant strains (Fig. 5C). Moreover, the *tsa1*Δ mutant and other strains did not exhibit DSB formation outside of the RFB site (Fig. 5B).

A previous study demonstrated that not only DSB formation but also end resection induces homologous recombination-mediated rDNA copy number changes (Sasaki and Kobayashi, 2017; 2023a; Sasaki and Kobayashi, 2023b). Resected DSBs that appeared below DSB fragments were observed in the *sir2*Δ mutant (Sasaki and Kobayashi, 2023a). However, they were barely detected in the WT and *tsa1*Δ mutant (Fig. 5B). Notably, we did not use cells synchronized in S phase, and the signals for resected DSBs were too low in our samples; thus, we could not accurately quantify a slight increase in the frequency of DSBs, if any were present. Transcription from E-pro inhibits the association of cohesin complexes with cohesin-associated regions located between the origin of DNA replication and 5S rDNA, and its inhibition results in the induction of rDNA copy number changes during DSB repair (Laloraya et al., 2000; Kobayashi *et al*., 2004; Kobayashi and Ganley, 2005). Thus, we examined whether Tsa1 affects transcription from E-pro. We extracted total RNA from the WT and *tsa1*Δ strains, separated RNA by agarose gel electrophoresis under denaturing conditions and detected the transcripts from E-pro by Northern blotting with strand-specific probes (Fig. 5A). The *sir2*Δ mutant, in which transcription from E-pro is derepressed (Kobayashi and Ganley, 2005), was analyzed in parallel. An IGS1-F product ranging from ∼800 to ∼1,200 nt in size was detected in the *sir2*Δ mutant but was barely detected in the WT cells (Fig. 5D), consistent with previous findings (Houseley et al., 2007; Hosoyamada *et al*., 2019). The level of IGS1-F transcripts in the *tsa1*Δ mutant was comparable to that in WT cells (Fig. 5E).

**Figure 5.**
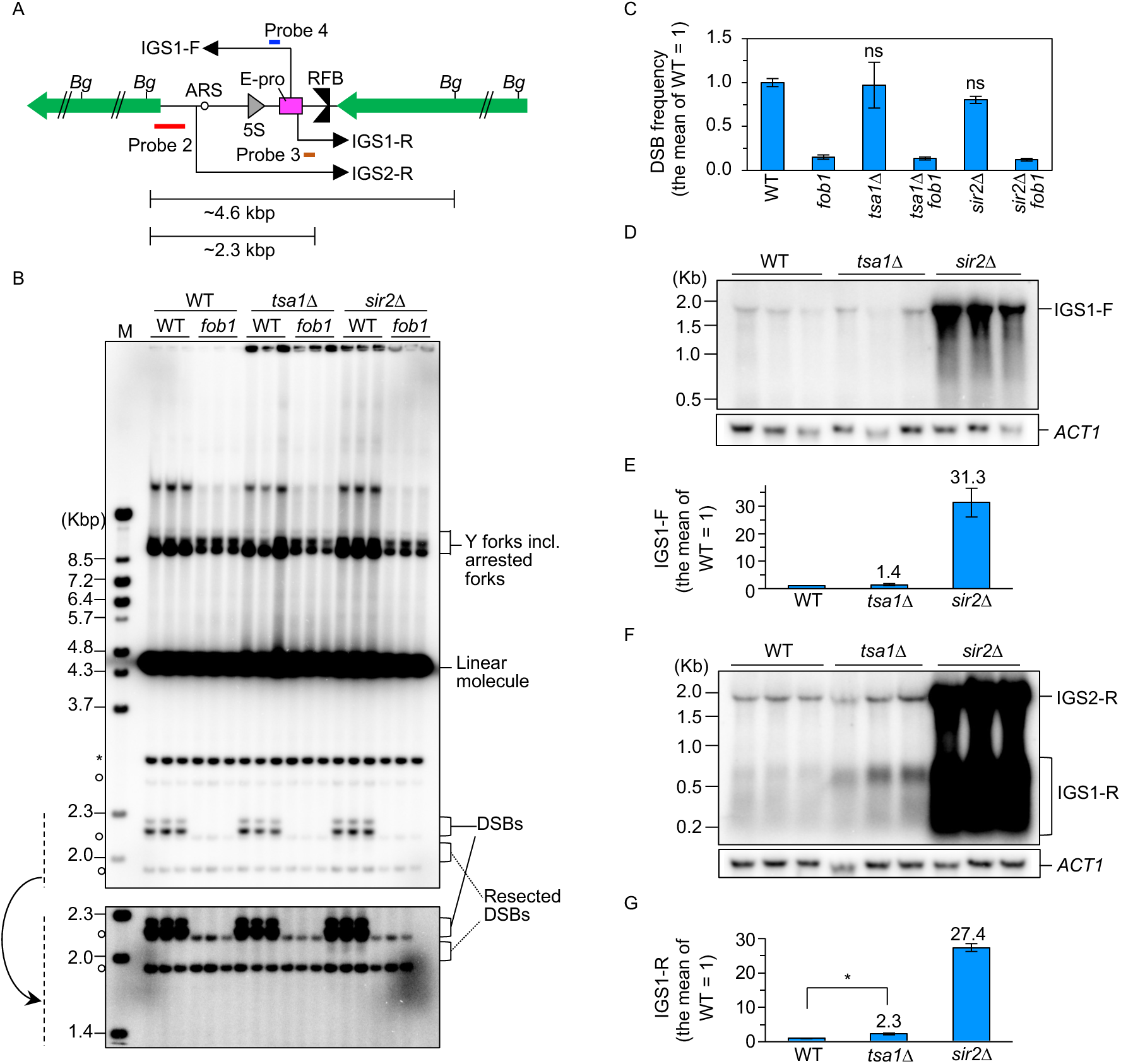
Transcription of IGS1-R from E-pro is stimulated in the *tsa1*Δ mutant. **(A)** Restriction map and positions of probes used for DSB analyses and Northern blotting. The restriction sites for BglII (Bg), the positions of probe 2 used for DSB analysis and probes 3 and 4 used for noncoding RNA detection by Northern blot analyses are indicated. **(B)** DSB assay. Genomic DNA was isolated from three clones of the indicated strains, digested with BglII, separated by size, and analyzed by Southern blotting with rDNA probe 2, as shown in (A). Asterisk and open circles on the left-hand side of the image indicate terminal fragments containing the telomere-proximal rDNA repeat and nonspecific background, respectively. Bands corresponding to DNA replication intermediates that include arrested forks in the *FOB1* background, linear fragments, DSBs, and resected DSBs are indicated on the right side of the image. The lower panel shows a more exposed contrast image of the phosphorimager signal around DSBs marked by dashed lines. **(C)** Quantitation of DSBs. The DSB frequency was determined by quantifying the signal of DSBs in (B) relative to that of replication intermediates, including arrested forks, which was normalized to the average of WT clones (bars show the mean ± s.e.m.). Multiple comparisons were performed by one-way ANOVA, followed by Tukey’s multiple comparisons test; ns, no significant difference (p > 0.05). **(D, F)** Detection of noncoding RNAs transcribed from E-pro. Total RNA from the indicated strains was separated by size on formaldehyde-agarose, followed by Northern blotting. In (D), IGS1-F was detected with probe 4 as indicated in (A). In (F), IGS1-R and IGS2-R were detected with probe 3 as indicated in (A). The membranes were reprobed for *ACT1* transcripts as a loading control. **(E, G)** Quantitation of noncoding RNA. Signals corresponding to IGS1-F (E) and IGS1-R transcripts (G) were quantified relative to ACT1, which was normalized to the average of WT clones. The bars show the means ± s.e.m. The transcript levels were compared between the WT and the *tsa1*Δ mutant strains by two-sided Student’s t tests; *, statistically significant difference (p < 0.05); ns, no significant difference (p > 0.05).

Toward the RFB, IGS1-R transcripts of ∼300–650 nt were observed in the *sir2*Δ mutant at a level ∼30-fold greater than that in WT cells (Fig. 5F) (Houseley *et al*., 2007; Hosoyamada *et al*., 2019). In the *tsa1*Δ mutants, IGS1-R transcripts were elevated by ∼2.3-fold compared with those in the WT, but their level was ∼10-fold lower than that in the *sir2*Δ mutant (Fig. 5G). The *sir2*Δ mutant showed an increase in the level of both IGS1-F and IGS1-R, but the *tsa1*Δ mutant only exhibited enhanced IGS1-R transcription (Fig. 5D–5G). The direction-specific impact of Tsa1 is currently unknown. Nonetheless, these findings imply that the function of Tsa1 negatively regulates IGS1-R transcription, which might stabilize cohesin associations and prevent rDNA copy number changes during DSB repair.

### Tsa1 suppresses SSB formation

ROS introduces oxidative damage to DNA bases and sugar phosphates, leading to the formation of 8-oxo-7,8-dihydroguanine (8-oxoG). 8-oxoG is primarily repaired by base excision repair, in which the modification is removed by the DNA glycosylase Ogg1, and the phosphodiester backbone is then excised by Ape1. The abortion of subsequent repair processes leads to the accumulation of SSBs (Slupphaug, 2003). In our DSB analysis, we did not detect DSBs outside the RFB in the WT*, sir2*Δ, or *tsa1*Δ strains (Fig. 5B). Thus, we examined whether the *tsa1*Δ mutant accumulated SSBs. To this end, the same genomic DNA samples that were analyzed for DSB detection were digested with Bgl II and separated by agarose gel electrophoresis under denaturing alkaline conditions, followed by Southern blotting (Fig. 6). In the WT, *tsa1*Δ, and *sir2*Δ strains, strong signals were detected at ∼ 2.3 kb, which corresponded to the template strand for DNA replication as well as newly synthesized strands made up of DSB fragments at the RFB (Fig. 6B). The *tsa1*Δ mutant additionally displayed a signal at ∼1.5 kb, but this signal was not detected in the WT, *fob1, sir2*Δ, or *sir2*Δ *fob1* mutant (Fig. 6B). Furthermore, this signal was also detected in the *tsa1*Δ *fob1* mutant. We did not detect bands at ∼1.5 kbp in our DSB analysis (Fig. 5B). Therefore, the *tsa1*Δ mutant experienced SSBs near 5S rDNA in an Fob1-independent manner, potentially inducing rDNA instability.

**Figure 6.**
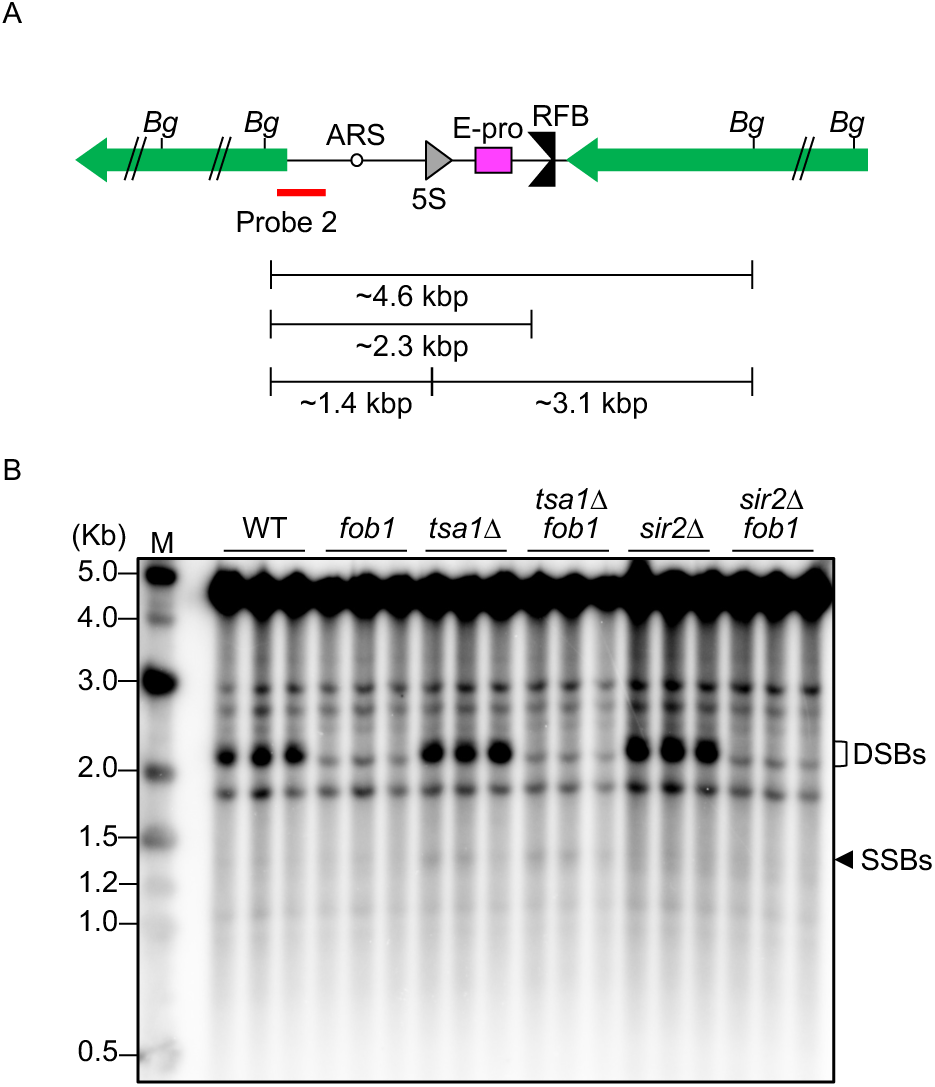
Tsa1 suppresses SSB formation in the rDNA. **(A)** Restriction map and positions of a probe used for SSB analyses. **(B)** SSB assay. Genomic DNA was isolated from three clones of the indicated strains, digested with BglII, and separated by agarose gel electrophoresis under denaturing conditions, followed by Southern blotting with rDNA probe 2, as shown in (A). Bands corresponding to DSB fragments at the RFB and SSBs are indicated.

### Tsa1-mediated rDNA stabilization contributes to lifespan extension

To assess which functions of Tsa1 are important for the maintenance of rDNA stability, we examined the effects of mutations in *TSA1* that compromise the ability of Tsa1 to affect rDNA stability. Cys48 and Cys171 of Tsa1 are critical for peroxidase activity *in vitro* (Fig. 1B) (Chae et al., 1994). Smearing of the chr XII band in the *tsa1*Δ mutant was suppressed by the introduction of a plasmid carrying *TSA1*, but it was not suppressed by the expression of *tsa1^C48S^*, *tsa1^C171S^*, or *tsa1^C48SC171S^* (Fig. 7A), indicating that both Cys48 and Cys171 are essential for the maintenance of rDNA stability. Phe44 and Tyr78 are important for the decamer formation of Tsa1 (Fig. 1B) (Loberg et al., 2019). The expression of *tsa1^F44A^*, *tsa1^Y78A^*, or *tsa1^F44AY78A^* did not suppress rDNA instability in the *tsa1*Δ mutant. Tsa1 can be hyperoxidized to function as a nonperoxidase, which needs to be reduced by the sulfiredoxin Srx1 to restore peroxidase activity (Fig. 1B) (Hanzen *et al*., 2016). The *srx1*Δ mutant itself showed homogeneous chr XII bands, similar to those in WT cells (Fig. 7B). Deletion of *SRX1* from the *tsa1*Δ mutant did not exacerbate rDNA instability in the *tsa1*Δ mutant (Fig. 7B). Therefore, peroxidase activity and decameric formation, but not Srx1, are important for rDNA stabilization.

**Figure 7.**
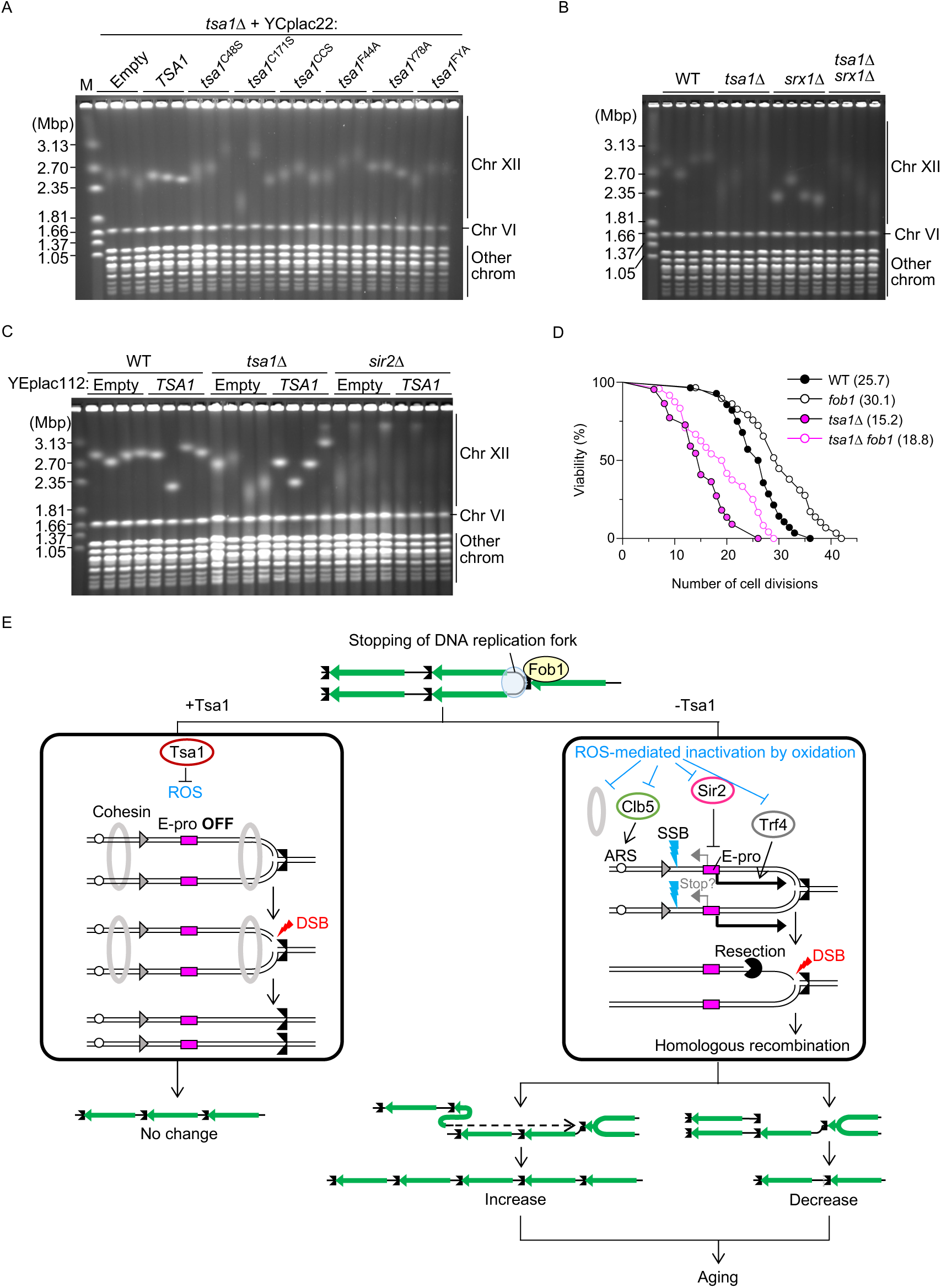
The Tsa1-mediated extension of the replicative lifespan depends on *FOB1*. **(A–C)** PFGE analysis of the size heterogeneity of chr XII. DNA was extracted from the indicated strains and separated by PFGE. DNA was stained with ethidium bromide. M indicates *H. wingei* chromosomal DNA markers. In (A), the *tsa1*Δ mutant was transformed with the YCplac22 vector, which is maintained in a cell at a single copy, and YCp22, which carries different *tsa1* alleles. In (C), the WT, *tsa1*Δ and *sir2*Δ strains were transformed with the mutlicopy vector YEplac112 or YCplac112 carrying *TSA1*. **(D)** Lifespan analysis of the indicated strains. Lifespan was measured by counting the number of cells that were produced from mother cells before death. The numbers of cells analyzed were 28 (WT), 29 (*fob1*), 22 (*tsa1*Δ) and 24 (*tsa1*Δ *fob1*Δ). The numbers in parentheses are the average lifespan. Kaplan-Meier survival curves were generated. For the statistical analysis, a log-rank test was performed: *P*=0.001 for WT versus *fob1*; *P*<0.0001 for WT versus *tsa1*Δ; *P*=0.03 for *tsa1*Δ versus *tsa1*Δ *fob1*; *P*<0.0001 for *fob1* versus *tsa1*Δ *fob1*. **(E)** Model of Tsa1-mediated rDNA stabilization. See the text for details.

Overexpression of *TSA1* does not reduce the mutation rate but extends the replicative lifespan through the redox switch of Tsa1 to disaggregate misfolded proteins (Hanzen *et al*., 2016). We analyzed whether *TSA1* overexpression also affects rDNA stability. To this end, we constructed a multicopy vector carrying *TSA1*, introduced it into WT, *tsa1*Δ, and *sir2*Δ mutants, and assessed rDNA stability by PFGE. WT strains transformed with an empty vector already showed sharp chr XII bands, so the stabilizing effects of Tsa1 overexpression, if present at all, could not be assessed (Fig. 7C). rDNA instability in the *tsa1*Δ mutant was suppressed by overexpression of *TSA1,* but we were unable to assess whether its overexpression resulted in more stable rDNA than that in the WT. The *sir2*Δ mutant showed smeared chr XII, but this smearing was not suppressed by overexpression of *TSA1* (Fig. 7C). Thus, the increased dosage of Tsa1 did not have a substantial impact on rDNA stability.

Finally, we examined whether suppression of rDNA instability by a *fob1* mutation prolongs the lifespan of the *tsa1*Δ mutant by counting the number of daughter cells produced from mother cells. Compared with that of the WT, the replicative lifespan of the *fob1* mutant was 20% longer, which is consistent with previous findings (Fig. 7D) (Defossez *et al*., 1999; Takeuchi *et al*., 2003). Moreover, the lifespan of the *tsa1*Δ mutant was shortened by ∼40% compared with that of the WT cells, consistent with previous findings (Molin *et al*., 2011; Nystrom *et al*., 2012; Roger *et al*., 2020). The lifespan of the *tsa1*Δ mutant was extended by 30% by the introduction of a f*ob1* mutation, although the lifespan of the *tsa1*Δ *fob1* mutant was still shorter than that of the WT (Fig. 7D, 7E). These lifespan patterns correlated well with the degree of rDNA instability of the *tsa1*Δ mutant in the *FOB1* and *fob1* backgrounds (Fig. 3A). Therefore, in addition to the previously known functions of Tsa1, such as molecular chaperone and redox signaling, Tsa1 delays cellular senescence through its ability to maintain rDNA stability.

## DISCUSSION

Genome instability and ROS impact aging in various organisms. In budding yeast, instability of the rDNA locus is one of the hallmarks of cellular aging. In this study, we aimed to determine whether the rDNA locus, which occupies nearly 10% of the entire budding yeast genome, is a target of ROS-mediated regulation of cellular aging. We screened 54 genes that control the production of ROS for their involvement in the maintenance of rDNA stability (Fig. 1). We have shown that *TSA1*, which encodes a major Prx in budding yeast, has an important function in the maintenance of rDNA stability.

Tsa1 suppresses the formation of recombination intermediates and rDNA instability in a manner dependent mainly on Fob1 (Fig. 3 and 4E, 4F). Analysis of rDNA replication intermediates by 2D gel analysis revealed a reduced frequency of replication initiation in the rDNA in the *tsa1*Δ mutant by ∼20% compared with that in the WT (Fig. 4B, 4C). Once DNA replication is initiated in the rDNA, more than 90% of replication forks that progress toward the 3’ end of the 35S rDNA are stalled at the RFB site (Brewer *et al*., 1992). Considering that fewer origins fired in the *tsa1*Δ mutant, fewer forks would be arrested at the RFB site in this mutant. However, the number of arrested forks was comparable between WT and *tsa1*Δ cells (Fig. 4D). These findings indicate that while fewer forks are stalled at the RFB in the *tsa1*Δ mutant, these forks tend to be more persistently arrested. We previously demonstrated that cells lacking the S-phase cyclin Clb5 exhibit more persistent arrest of forks at the RFB due to reduced replication initiation and the induction of HR-dependent rDNA instability without altering the level of DSBs (Goto *et al*., 2021). Similar phenotypes were observed for the *tsa1*Δ mutant, while the replication initiation frequency decreased by only ∼20% in the *tsa1*Δ mutant (Fig. 4, 5B, 5C) compared to ∼50% in the *clb5*Δ mutant (Goto *et al*., 2021). Thus, persistently arrested forks at the RFB may contribute to HR-dependent rDNA copy number changes, independent of DSBs that were observed in the fission yeast replication termination sequence RTS1 (Lambert et al., 2005; Mizuno et al., 2013).

DSBs are thought to be major triggers of genome rearrangements, especially when they are formed in repetitive sequences (Sasaki et al., 2010). Budding yeast cells maintain rDNA stability during the repair of DSBs formed at the RFB (Fig. 1A) (Sasaki and Kobayashi, 2023a). First, DSB end resection is normally suppressed for DSBs at the RFB, but when this suppression is relieved, DSBs are repaired by HR, which tends to induce rDNA copy number changes (Sasaki and Kobayashi, 2017; Sasaki and Kobayashi, 2023b). Because we used log-phase cells where the proportion of S-phase cells was small in this study, we were not able to assess the effect of Tsa1 deficiency on DSB end resection (Fig. 5B). Second, the maintenance of rDNA copy number during DSB repair is also ensured by alignment of the rDNA copies in an equal position by cohesin complexes (Kobayashi *et al*., 2004). Transcription from E-pro not only inhibits cohesin association but also relieves the suppression of DSB end resection (Kobayashi and Ganley, 2005; Sasaki and Kobayashi, 2023b). Tsa1 deficiency led to an increase in the IGS1-R transcript level (Fig. 5D–5G, 7E right). The level of transcripts that remain in cells is influenced by the balance between the synthesis and degradation of transcripts. When Sir2 is absent, E-pro is derepressed, and the transcription of noncoding RNA in both directions is increased (Kobayashi and Ganley, 2005). The absence of the Ccr4/Pop2/Not mRNA deadenylase complex leads to the accumulation of both IGS1-F and IGS1-R transcripts (Hosoyamada *et al*., 2019), while the absence of the RNA poly(A) polymerase Trf4 causes defective degradation of IGS1-R transcripts (Houseley *et al*., 2007). In the *tsa1*Δ mutant, the functions of any of these factors may be compromised by their oxidation by ROS accumulation, resulting in defective regulation of transcription from E-pro, inhibition of cohesin binding and rDNA instability (Fig. 7E, right). The mechanisms underlying IGS1-R accumulation need to be examined in future studies.

ROS can generate various kinds of DNA modifications, such as 8-oxoG. This damage is repaired by base excision repair, in which the modification is removed by the DNA glycosylase Ogg1, and the phosphodiester backbone is then excised by Ape1. The abortion of subsequent repair processes leads to the accumulation of SSBs (Slupphaug, 2003). Thus, when Tsa1 was absent, it is possible that ROS-dependent DNA damage frequently occurred near the 5S rDNA for reasons that remain to be studied in future studies, its repair was frequently aborted, and SSBs remained unrepaired (Fig. 5F, 7E right). Although highly speculative, these SSBs may inhibit the progression of the transcription machinery that progresses from E-pro to the 5S rDNA.

Tsa1 is known to function as a peroxidase, molecular chaperone, and redox-dependent modulator. The peroxidase activity of Tsa1 depends on oxidation‒reduction sites at Cys48 and Cys171 as well as at Phe44 and Tyr 78, which are crucial for decameric formation (Wood *et al*., 2003). All these residues were important for the maintenance of rDNA stability (Fig. 7A), indicating that the peroxidase function of Tsa1 most likely contributes to rDNA stabilization. It remains unclear why Srx1 is not important for the maintenance of rDNA stability (Fig. 7B), but we speculate that under physiological conditions without any exogenous H_2_O_2_, Tsa1 is rarely hyperoxidized or the level of hyperoxidized Tsa1 is below the threshold to destabilize rDNA; thus, cells do not require Srx1. The molecular chaperone activity of Tsa1 depends on structural changes to higher molecular weight forms, but this effect depends on Cys48 of Tsa1 and Srx1 but not Cys171 of Tsa1 (Jang *et al*., 2004). Our finding that Cys171 was also important for rDNA stability indicates that the contribution of the molecular chaperone activity of Tsa1 to rDNA stabilization is most likely minimal. For redox-dependent target modulation, both Cys171 and Cys48 are necessary for the activation of other proteins (Tachibana et al., 2009). Thus, this nonperoxidase function may be involved in promoting rDNA stability. Yap1 is one such target protein (Tachibana *et al*., 2009), but it is not required for the maintenance of rDNA stability (Fig. 2A). Factors that function to maintain rDNA stability could be targets of Tsa1-mediated oxidization, such as Sir2, the Pop2/Ccr4/NOT complex and Trf4, which regulate transcription from E-pro; Clb5, which promotes efficient replication initiation; cohesin complexes; and factors involved in base excision repair (Fig. 7E, right). This possibility needs to be explored in future studies.

Previous studies have demonstrated that Tsa1 delays aging through its chaperone function to facilitate the clearance of H_2_O_2_-mediated protein aggregates and its ability to modify protein kinase A in the nutrient signaling pathway (Hanzen *et al*., 2016; Roger *et al*., 2020). Furthermore, because the absence of Tsa1 causes a restricted lifespan in the *sir2*Δ *fob1*Δ background, it was suggested that the Tsa1-mediated delay of aging is independent of its ability to regulate ERCs (Hanzen *et al*., 2016). This is supported by our findings that ERCs accumulate less in the *tsa1*Δ mutant than in the *sir2*Δ mutant (Figure 2E and 3B).

In this study, we demonstrated that the stability of rDNA was lower in the *tsa1*Δ mutant than in the WT strain (Fig. 3). Furthermore, rDNA stability was greater in the *tsa1*Δ *fob1* mutant than in the *tsa1*Δ mutant but not in the WT cells (Fig. 3). In good agreement with the rDNA instability, the *tsa1*Δ *fob1* mutant had a shorter lifespan than did the *fob1* mutant, and the lifespan of the *tsa1*Δ *fob1* mutant did not improve to the lifespan of the WT cells (Fig. 7D). Therefore, in addition to its previously known function, the Tsa1-mediated maintenance of rDNA stability also contributes to lifespan extension (Fig. 7E).

## AUTHOR CONTRIBUTIONS

J.O., M.S. and T.K. designed the experiments, analyzed the data and wrote the paper. J.O. performed the experiments.

## ACKNOWLEDGEMENTS

We thank members of the Kobayashi laboratory for discussions. This work was supported by Grants-in-Aid for Scientific Research (20H05382 and 18H04709 to M.S. and 17H01443 and 21H04761 to T.K.), the JST FOREST Program (grant number JPMJFR214P to M.S.), and JST AMED (JP21gm1110010 to T.K.). JST CREST (grant number JPMJCR19S3 to T.K.).

## DECLARATION OF INTERESTS

The authors declare no competing interests.

## MATERIALS AND METHODS

### Yeast Strains and Culture Methods

The yeast strains used in this study are derivatives of W303 (*MATa ade2-1 ura3-1 his3-11, 15 trp1-1 leu2-3,112 can1-100*) and are listed in Table 1. Gene deletion was performed by standard one-step gene replacement methods, and genotypes were confirmed by PCR-based genotyping. Haploid mutant strains were constructed by gene deletion from the WT haploid strain. Alternatively, diploids heterozygous for gene deletions of interest were first constructed and subsequently subjected to sporulation, followed by the isolation of haploid clones by tetrad dissection. Prior to use, the yeast strains were streaked onto YPD plates (1% w/v yeast extract, 2% w/v peptone, 2% w/v glucose, and 2% w/v agar). Yeast strains were inoculated into YPD liquid media (1% w/v yeast extract, 2% w/v peptone, 2% w/v glucose), unless otherwise noted, and were grown at 30°C.

For PFGE and ERC analyses, single colonies of yeast strains were inoculated into YPD liquid media and shaken overnight until the cultures reached the stationary phase. The cells were collected (5 × 10^7^ cells/plug) and washed twice with 50 mM EDTA (pH 7.5). For two-dimensional (2D) agarose gel electrophoresis and DSB analyses, overnight cultures were diluted in 50 mL of YPD medium to an OD_600_ = 0.1 and grown to an OD_600_ = 0.4. The cultures were immediately treated with 1/1,000 vol of 10% sodium azide, collected (5 × 10^7^ cells/plug) by centrifugation for 2 min at 2,400 × *g* at 4°C and washed twice with 50 mM EDTA (pH 7.5). To prepare RNA, overnight cultures were diluted in 15 mL of YPDA medium (YPD containing 40 μg/mL adenine sulfate) to an OD_600_ = 0.1 and collected when the cultures reached an OD_600_ of 0.8. SC-trp

### Plasmid construction

YCplac22 and YEplac22 plasmids that carry *TSA1* were constructed by the Gibson assembly method as follows. The genomic regions containing the *TSA1* open reading frame and ∼500 bp upstream and downstream were amplified by PCR from the *S. cerevisiae* WT strain on the W303 background. Linearized YCplac22 and YEplac22 were generated by PCR from YCplac22 and YEplac22. These PCR fragments had short homology at each end and were fused using the gibsom assembly according to the manufacturer’s instructions. YCplac22 plasmids that carry *tsa1* mutant alleles were constructed using the PrimeSTAR Mutagenesis Basal Kit (TaKaRa) according to the manufacturer’s instructions. Briefly, primers were designed to have 15 nucleotides of overlap at their 5’ ends and to have mutations that change the amino acid residue of interest into different residues. The primers used were as follows. These primers were used to perform PCR using YCplac22-TSA1 as a template. The resulting PCR products were subsequently transformed into *E. coli* DH5alpha. Plasmid DNA was sequenced to confirm that the mutations were properly introduced. The sequences of primers used are available upon request.

### Genomic DNA preparation

For PFGE, ERC, DSB, and 2D gel analyses, genomic DNA was prepared in low melting temperature agarose plugs as described previously (Sasaki and Kobayashi, 2017; 2021). Briefly, collected cells were resuspended in 50 mM EDTA (pH 7.5; 33 μL of cells per 5 × 10^7^ cells) and incubated at 42°C. For each plug, 33 μL of cell suspension was mixed with 66 μL of solution 1 (0.83% low-melting-point agarose SeaPlaque GTG (Lonza), 170 mM sorbitol, 17 mM sodium citrate, 10 mM EDTA (pH 7.5), 0.85% β-mercaptoethanol, and 0.17 mg/mL Zymolyase 100 T (Nacalai)), poured into a plug mold (Bio-Rad) and placed at 4°C for agarose solidification. Plugs were transferred to a 2-mL tube containing solution 2 (450 mM EDTA pH 7.5, 10 mM Tris-HCl pH 7.5, 7.5% β-mercaptoethanol and 10 μg/mL RNaseA (Macherey-Nagel)) and incubated for 1 h at 37°C. Plugs were then incubated overnight at 50°C in solution 3 (250 mM EDTA pH 7.5, 10 mM Tris-HCl pH 7.5, 1% sodium dodecyl sulfate (SDS) and 1 mg/mL proteinase K (Nacalai)). The plugs were subsequently washed four times with 50 mM EDTA (pH 7.5) and stored at 4°C in 50 mM EDTA (pH 7.5).

### PFGE

PFGE was performed as described previously (Sasaki and Kobayashi, 2017; 2021). Briefly, one-third of a plug was placed on a tooth of the comb, including a piece with *Hansenula wingei* chromosomal DNA markers (Bio-Rad). The comb was placed in a gel tray, and 1.0% agarose solution (Pulsed Field Certified Agarose, Bio-Rad) in 0.5× TBE (44.5 mM Tris base, 44.5 mM boric acid and 1 mM EDTA, pH 8.0). PFGE was performed on a Bio-Rad CHEF DR-III system in 2.2 L of 0.5× TBE under the following conditions: 3.0 V/cm for 68 h at 14°C, 120°C, an initial switch time of 300 s, and a final switch time of 900 s. After electrophoresis, the DNA was stained with 0.5 μg/mL ethidium bromide (EtBr) for 30 min, washed with dH_2_O for 30 min, and then photographed.

### Southern blotting

#### Agarose gel electrophoresis

##### ERC assay

The ERC assay was performed as described previously (Goto *et al*., 2021; Sasaki and Kobayashi, 2021). Half of the agarose plug was placed on the tooth of the comb. After the comb was set in the gel tray (15 × 25 cm), 300 ml of 0.4% agarose (SeaKem LE Agarose, Lonza) in 1× TAE (40 mM Tris base, 20 mM acetic acid, and 1 mM EDTA pH 8.0) was poured into the tray and allowed to set, after which 500 ng of lambda HindIII DNA marker was applied to an empty lane. The electrophoresis was performed on a subcell GT electrophoresis system (Bio-Rad) in 1.5 L of 1× TAE at 1.0 V/cm for ∼48 h at 4°C with buffer circulation. The buffer was changed every ∼24 h. DNA was stained with 0.5 μg/mL EtBr for 30 min and then photographed.

##### 2D gel electrophoresis

2D gel electrophoresis was performed as described previously with slight modifications (Goto *et al*., 2021). One-half of an agarose plug was placed in a 2-mL tube. The plug was equilibrated twice in 1 mL of 1× M buffer (TaKaRa) by rotating the tube for 30 min at room temperature. After discarding the buffer, the plug was incubated in 160 μL of 1x M buffer containing 160 units of *Nhe* I (TaKaRa) overnight at 37°C. The plug was placed on the tooth of the comb. The comb was placed in a gel tray (15 × 25 cm), 0.4% agarose solution in 1× TBE was added, and the gel was allowed to solidify. A total of 600 ng of lambda HindIII DNA markers was applied to the empty lane. The electrophoresis was performed on a subcell GT electrophoresis system (Bio-Rad) in 1.5 L of 1× TBE at 1.32 V/cm for 14 h at room temperature with buffer circulation.

After electrophoresis, the DNA was stained with 0.3 μg/mL EtBr for 30 min and then photographed. Gel slices containing DNA ranging from 4.7 to 9.4 kb were excised, rotated 90°, and placed onto the gel tray used for the second agarose gel electrophoresis, for which 1.2% agarose solution in 1x TBE containing 0.3 μg/mL of EtBr was poured into the tray. Second-dimensional electrophoresis was performed on a subcell GT electrophoresis system (Bio-Rad) in 1.5 L of 1× TBE containing 0.3 μg/mL EtBr at 6.0 V/cm for 6 h at 4°C with buffer circulation. After electrophoresis, the DNA was photographed.

##### DSB assay

The DSB assay was performed as described previously (Sasaki and Kobayashi, 2017; 2021). One-third of an agarose plug was cut and placed in a 2-mL tube. The plug was equilibrated four times in 1 mL of 1× TE (10 mM Tris base, pH 7.5, and 1 mM EDTA, pH 8.0) by rotating the tube for 15 min at room temperature. The plug was equilibrated twice in 1 mL of 1× NEBuffer 3.1 (New England Biolabs) by rotating the tube for 30 min at room temperature. After discarding the buffer, the plug was incubated in 160 μL of 1× NEBuffer 3.1 buffer containing 160 units of *Bgl* II (New England Biolabs) overnight at 37°C. The plug was placed on a tooth of the comb, which was set into the gel tray (15 × 25 cm), into which 0.7% agarose solution in 1x TBE was poured; after setting the gel, 600 ng of lambda *Bst* EⅡ DNA markers were applied to an empty lane. The electrophoresis was performed on a subcell GT electrophoresis system (Bio-Rad) in 1.5 L of 1× TBE at 2.0 V/cm for 22 h at room temperature with buffer circulation. After electrophoresis, the DNA was stained with 0.5 μg/mL EtBr for 30 min and then photographed.

##### SSB assay

One-third of an agarose plug was washed and digested with BglII in the same way as the plugs subjected to the DSB assay. The BglII reactions were discarded. The plugs were washed with 1 ml of dH_2_O by rotating the tube for 10 min at room temperature, after which the dH_2_O was discarded. This process was repeated one more time. The plugs were equilibrated in 5× alkaline running buffer (0.25 M NaOH, 5 mM EDTA [pH 8.0]) by rotating for 15 min at room temperature, after which the buffer was discarded. This process was repeated one more time. The buffer was discarded. Plugs were equilibrated twice in 1× alkaline running buffer (0.05 M NaOH, 1 mM EDTA [pH 8.0]) by rotating for 15 min at room temperature, after which the buffer was discarded. This process was repeated one more time. The plug was placed on the tooth of the comb, which was set into the gel tray (15 × 20 cm) in a cold room.

The agarose solution was prepared by dissolving 2.5 g of Seakem LE agarose (Lonza) in 225 ml of Milli-Q water in a microwave, followed by cooling to 50°C. Then, 25 ml of 10x alkaline buffer (0.5 M NaOH, 10 mM EDTA [pH 8.0]) was added to the agarose solution, mixed, poured into the gel tray in a cold room, and left until the gel solidified. The comb was removed, and 1.45 L of 1× alakaline buffer was added. DNA markers of 100 bp ladder (NEB) and 1 kb ladder (NEB) were mixed with 1× alkaline buffer, 0.1% bromophenol blue and 5% glycerol, and the mixture was heat-denatured for 5 min at 70°C, followed by rapid chilling on ice. After circulating the buffer for 15 minutes with a peristaltic pump, 10 µl of DNA marker was added, and the gel was run overnight at 4°C at 50 V. After electrophoresis, the gel was immersed in sterile water, followed by agitation for 45 minutes in neutralization buffer (0.5 M Tris-HCl, 1 M NaCl). This agitation was repeated twice, followed by agitation for 45 minutes in 0.5 µg/ml ethidium bromide. Gel photos were captured via UV exposure.

#### DNA transfer

After agarose gel electrophoresis, the gel was gently mixed in 500 mL of 0.25 N HCl for 30 min, 500 mL of denaturation solution (0.5 N NaOH, 1.5 M NaCl) for 30 min, and 500 mL of transfer buffer (0.25 N NaOH, 1.5 M NaCl) for 30 min. DNA was transferred to Hybond-XL (GE Healthcare) by the standard capillary transfer method with transfer buffer. DNA was fixed to the membrane by soaking the membrane in 300 mL of freshly prepared 0.4 N NaOH for 10 min with gentle shaking, followed by rinsing the membrane with 2x SSC for 10 min. The membrane was subsequently dried and stored at 4°C.

#### Probe preparation

Probes were prepared as described previously (Sasaki and Kobayashi, 2017; 2021). Double-stranded DNA fragments were amplified by PCR. Probe 1 used for ERC and 2D analyses was amplified with the primers 5’-CATTTCCTATAGTTAACAGGACATGCC and 5’-AATTCGCACTATCCAGCTGCACTC, and probe 2 used for DSB analysis was amplified with the primers 5’-ACGAACGACAAGCCTACTCG and 5’-AAAAGGTGCGGAAATGGCTG. A portion of the PCR products was gel-purified and seeded for a second round of PCR with the same primers. The PCR products were gel-purified, and 50 ng was used for random priming reactions in the presence of the radiolabeled nucleotide [α-^32^P]-dCTP (3,000 Ci/mmol, 10 mCi/ml, Perkin Elmer) using a random primer DNA labeling kit (TaKaRa) according to the manufacturer’s instructions. Unincorporated nucleotides were removed using ProbeQuant G-50 Micro Columns (GE Healthcare). The radiolabeled probes were heat denatured for 5 min at 100°C immediately prior to hybridization to the membrane.

#### Hybridization

Southern hybridization was performed as described previously (Sasaki and Kobayashi, 2017; 2021). The membrane was prewetted with 0.5 M phosphate buffer, pH 7.2, and prehybridized for 1 h at 65°C with 25 mL of hybridization buffer (1% bovine serum albumin, 0.5 M phosphate buffer, pH 7.2; 7% SDS; 1 mM EDTA, pH 8.0). After the buffer was discarded, the membrane was hybridized with 25 mL of hybridization buffer and heat-denatured probe overnight at 65°C. The membrane was washed four times for 15 min at 65°C with wash buffer (40 mM phosphate buffer pH 7.2, 1% SDS, 1 mM EDTA pH 8.0) and exposed to a phosphor screen.

#### Image analysis

Membranes were exposed to a phosphor screen for several days to achieve a high signal-to-noise ratio for DNA molecules that are low in abundance, such as ERCs, replication intermediates in 2D assays, and arrested forks and DSBs. The radioactive signal was detected using a Typhoon FLA 7000 (GE Healthcare). In the ERC assays, the genomic rDNA signal was used for normalization of the ERC signals. Thus, the membranes were re-exposed to the phosphor screen for a short time before this signal reached saturation. ERC bands and genomic rDNA were quantified using FUJIFILM Multi Gauge version 2.0 software (Fujifilm) and scans from long and short exposures, respectively. The levels of ERCs were calculated by dividing the sum of the signal intensities of the ERCs by those of the genomic rDNA. When the levels of ERCs were compared between samples loaded on different gels, pieces of the plug excised from a specific DNA sample were loaded on different gels, blotted and hybridized. The signals of these plugs were used for normalization of samples on different gels. In 2D analyses, bubbles, Y arcs, RFB spots, and double Y spots were quantified using ImageJ (NIH). The RFB activity was calculated as the ratio between the signal for the spot with arrested forks and the sum of signals encompassing the total number of DNA replication intermediates. The signals of DSBs and arrested forks were quantified using FUJIFILM Multi Gauge version 2.0 software (Fujifilm). The frequency of DSBs was calculated by normalizing the DSB signal to that of the arrested forks.

### Yeast RNA preparation

RNA was prepared as described previously (Iida and Kobayashi, 2019), with slight modifications. The collected cells were resuspended in 400 μL of TES (10 mM Tris-HCl pH 7.5, 10 mM EDTA pH 7.5, 0.5% SDS) and 400 μl of acidic phenol by vortexing for approximately 10 s. The cells were incubated at 65°C for 1 h with occasional vortexing every 15 min. The cell suspensions were incubated for 5 min on ice and centrifuged at 20,000 × *g* for 10 min at 4°C. The aqueous phase was transferred to a new tube and mixed with an equal volume of acidic phenol by vortexing for 10 s. Samples were incubated for 5 min on ice and centrifuged at 20,000 × *g* for 10 min at 4°C. The aqueous phase was transferred to a new tube. Then, 1/10 vol. of 3 M sodium acetate (pH 5.3) and 2.5 vol. of 100% ethanol were added, and RNA was precipitated overnight at −20°C. After centrifugation at 20,000 × *g* for 10 min at 4°C, the RNA was washed with 70% ethanol. The RNA pellets were resuspended in 30 μL of dH_2_O and treated with 0.1% diethylpyrocarbonate (DEPC). The concentration of RNA was quantified using a NanoDrop ND-1000 spectrophotometer (Thermo Fisher Scientific). RNA samples were stored at −80°C.

### Northern blotting

Total RNA (18∼30 μg) was added to 7 μL of DEPC-treated dH_2_O and mixed with 17 μL of RNA sample buffer (396 μL of deionized formamide, 120 μL of 10x MOPS buffer [0.2 M MOPS, 50 mM sodium acetate pH 5.2, 10 mM EDTA pH 7.5 in DEPC-treated dH_2_O], and 162 μL of 37% formaldehyde). The samples were heated at 65°C for 20 min, followed by rapid chilling on ice. Six microliters of 6x Gel Loading Dye (B7025S, New England Biolabs) and 1.5 μL of 2.5 mg/ml EtBr were added to each sample. A 1% agarose gel was made by dissolving 1 g of agarose powder in 73 mL of DEPC-treated dH_2_O, and after cooling to 60°C, 17 mL of 37% formaldehyde and 10 mL of 10× MOPS buffer were added. The solution was poured into a gel tray (13 × 12 cm) and allowed to set; 10 μg of total RNA (10 μL) and 1.8 μg of DynaMarker RNA High markers were applied. Agarose gel electrophoresis was performed on a Mupid EX system (Takara) in 400 mL of 1× MOPS buffer at 20 V for 20 min and then at 100 V until the bromophenol blue dye had migrated approximately 2/3 of the gel. After electrophoresis, the gel was photographed, and RNA was transferred to Hybond-N+ (GE Healthcare) by standard capillary transfer.

Strand-specific probes were prepared from double-stranded DNA fragments, amplified by PCR and gel-purified. The PCR primers for probe 3 (against IGS1-F) were 5’-AGGGAAATGGAGGGAAGAGA and 5’-TCTTGGCTTCCTATGCTAAATCC; for probe 4 (against IGS1-R, IGS2-R), 5’-TCGCCAACCATTCCATATCT and 5’-CGATGAGGATGATAGTGTGTAAGA; and for detecting ACT1, 5’-CGAATTGAGAGTTGCCCCAG and 5’-CAAGGACAAAACGGCTTGGA. Strand-specific probes were then prepared by linear PCR in a final volume of 20 μL containing 0.2 mM dATP, 0.2 mM dTTP, 0.2 mM dGTP, 50 μL [α-^32^P]-dCTP (3,000 Ci/mmol, 10 mCi/ml, Perkin Elmer), 1.25 µL of ExTaq (TaKaRa), 1× ExTaq buffer, 50 ng of PCR product as a template, and 10 μM primer (5’-AGTTCCAGAGAGGCAGCGTA for probe 3, 5’-CATTATGCTCATTGGGTTGC for probe 4). PCR was initiated by a denaturation step at 94°C for 3 min, followed by 35 cycles of amplification (96°C for 20 s, 51°C for 20 s, and 72°C for 30 s), and a final step at 72°C for 3 min. Unincorporated nucleotides were removed using ProbeQuant G-50 Micro Columns (GE Healthcare). The radiolabeled probes were heat denatured by incubating for 5 min at 100°C immediately prior to hybridization to the membrane.

The membrane was incubated with 10 mL of ULTRAhyb Ultrasensitive Hybridization Buffer (Thermo Fisher) at 42°C for 1 h. The heat-denatured probe was incubated with the membrane overnight at 42°C. The membrane was rinsed twice with 2x SSC, washed for 15 min at 42°C twice with wash buffer 1 (2x SSC, 0.1% SDS), and washed for 15 min at 42°C twice with wash buffer 2 (0.1x SSC, 0.1% SDS). The membrane was exposed to phosphor screens for several days, and radioactive signals were detected using a Typhoon FLA 7000 (GE Healthcare). Probes were stripped by incubating the membrane with boiled 0.1% SDS by shaking for ∼30 min, rinsed with 2x SSC, and rehybridized with the ACT1 probe that was prepared as described above. Signals of IGS-F, IGS1-R, IGS2-R, and ACT1 were quantified using FUJIFILM Multi Gauge version 2.0 software (Fujifilm). The levels of IGS transcripts were normalized to the ACT-1 signal.

### Replicative lifespan analysis

The replicative lifespan was measured as previously described (Kennedy et al., 1994). In brief, the strains were streaked on a YPD plate and grown overnight at 30°C. The next day, the cells were streaked on a YPD plate at a low density and incubated at 30°C. Cells that emerged as small buds were transferred to other areas of the plate using the Singer micromanipulator MSM400, and the plate was incubated at 30°C. When these cells produced buds, the budded daughter cells were removed using a micromanipulator. When the experiments were paused, the plates were placed in an incubator at 10°C. These procedures were continued until the mother cell stopped producing more buds. When divisions were not observed in six observations, we judged that the mother cells died and stopped counting. The number of budded daughter cells was counted and designated as the replicative lifespan of each mother cell. A Kaplan‒Meier survival curve was generated for the replicative life span. Statistical significance was determined by the log-rank test, and significance was defined as *P*<0.05.

## Notes

### Competing Interest Statement

The authors have declared no competing interest.

## REFERENCES

Brewer, B.J., Lockshon, D., and Fangman, W.L. (1992). The arrest of replication forks in the rDNA of yeast occurs independently of transcription. Cell 71, 267–276. 10.1016/0092-8674(92)90355-g.

Burkhalter, M.D., and Sogo, J.M. (2004). rDNA enhancer affects replication initiation and mitotic recombination: Fob1 mediates nucleolytic processing independently of replication. Molecular cell 15, 409–421. 10.1016/j.molcel.2004.06.024.

Chae, H.Z., Uhm, T.B., and Rhee, S.G. (1994). Dimerization of thiol-specific antioxidant and the essential role of cysteine 47. Proceedings of the National Academy of Sciences of the United States of America 91, 7022–7026. 10.1073/pnas.91.15.7022.

Defossez, P.-A., Prusty, R., Kaeberlein, M., Lin, S.-J., Ferrigno, P., Silver, P.A., Keil, R.L., and Guarente, L. (1999). Elimination of Replication Block Protein Fob1 Extends the Life Span of Yeast Mother Cells. Molecular cell 3, 447–455. 10.1016/s1097-2765(00)80472-4.

Delaunay, A., Pflieger, D., Barrault, M.B., Vinh, J., and Toledano, M.B. (2002). A thiol peroxidase is an H2O2 receptor and redox-transducer in gene activation. Cell 111, 471–481. 10.1016/s0092-8674(02)01048-6.

Di Micco, R., Krizhanovsky, V., Baker, D., and d’Adda di Fagagna, F. (2021). Cellular senescence in ageing: from mechanisms to therapeutic opportunities. Nat Rev Mol Cell Biol 22, 75–95. 10.1038/s41580-020-00314-w.

Fang, J., and Beattie, D.S. (2003). External alternative NADH dehydrogenase of Saccharomyces cerevisiae: a potential source of superoxide. Free Radic Biol Med 34, 478–488. 10.1016/s0891-5849(02)01328-x.

Goto, M., Sasaki, M., and Kobayashi, T. (2021). The S-Phase Cyclin Clb5 Promotes rRNA Gene (rDNA) Stability by Maintaining Replication Initiation Efficiency in rDNA. Molecular and cellular biology 41. 10.1128/MCB.00324-20.

Hanzen, S., Vielfort, K., Yang, J., Roger, F., Andersson, V., Zamarbide-Fores, S., Andersson, R., Malm, L., Palais, G., Biteau, B., et al. (2016). Lifespan Control by Redox-Dependent Recruitment of Chaperones to Misfolded Proteins. Cell 166, 140–151. 10.1016/j.cell.2016.05.006.

Hosoyamada, S., Sasaki, M., and Kobayashi, T. (2019). The CCR4-NOT Complex Maintains Stability and Transcription of rRNA Genes by Repressing Antisense Transcripts. Molecular and cellular biology 40. 10.1128/MCB.00320-19.

Houseley, J., Kotovic, K., El Hage, A., and Tollervey, D. (2007). Trf4 targets ncRNAs from telomeric and rDNA spacer regions and functions in rDNA copy number control. EMBO J 26, 4996–5006. 10.1038/sj.emboj.7601921.

Huang, M.E., Rio, A.G., Nicolas, A., and Kolodner, R.D. (2003). A genomewide screen in Saccharomyces cerevisiae for genes that suppress the accumulation of mutations. Proceedings of the National Academy of Sciences of the United States of America 100, 11529–11534. 10.1073/pnas.2035018100.

Hughes, A.L., and Gottschling, D.E. (2012). An early age increase in vacuolar pH limits mitochondrial function and lifespan in yeast. Nature 492, 261–265. 10.1038/nature11654.

Iida, T., and Kobayashi, T. (2019). RNA Polymerase I Activators Count and Adjust Ribosomal RNA Gene Copy Number. Molecular cell 73, 645–654 e613. 10.1016/j.molcel.2018.11.029.

Iraqui, I., Kienda, G., Soeur, J., Faye, G., Baldacci, G., Kolodner, R.D., and Huang, M.E. (2009). Peroxiredoxin Tsa1 is the key peroxidase suppressing genome instability and protecting against cell death in Saccharomyces cerevisiae. PLoS Genet 5, e1000524. 10.1371/journal.pgen.1000524.

Irokawa, H., Tachibana, T., Watanabe, T., Matsuyama, Y., Motohashi, H., Ogasawara, A., Iwai, K., Naganuma, A., and Kuge, S. (2016). Redox-dependent Regulation of Gluconeogenesis by a Novel Mechanism Mediated by a Peroxidatic Cysteine of Peroxiredoxin. Sci Rep 6, 33536. 10.1038/srep33536.

Jang, H.H., Lee, K.O., Chi, Y.H., Jung, B.G., Park, S.K., Park, J.H., Lee, J.R., Lee, S.S., Moon, J.C., Yun, J.W., et al. (2004). Two enzymes in one; two yeast peroxiredoxins display oxidative stress-dependent switching from a peroxidase to a molecular chaperone function. Cell 117, 625–635. 10.1016/j.cell.2004.05.002.

Kaeberlein, M., McVey, M., and Guarente, L. (1999). The SIR2/3/4 complex and SIR2 alone promote longevity in Saccharomyces cerevisiae by two different mechanisms. Genes Dev 13, 2570–2580. 10.1101/gad.13.19.2570.

Kaniak-Golik, A., Kuberska, R., Dzierzbicki, P., and Sledziewska-Gojska, E. (2017). Activation of Dun1 in response to nuclear DNA instability accounts for the increase in mitochondrial point mutations in Rad27/FEN1 deficient S. cerevisiae. PLoS One 12, e0180153. 10.1371/journal.pone.0180153.

Kennedy, B.K., Austriaco, N.R., Jr., and Guarente, L. (1994). Daughter cells of Saccharomyces cerevisiae from old mothers display a reduced life span. J Cell Biol 127, 1985–1993. 10.1083/jcb.127.6.1985.

Kobayashi, T., and Ganley, A.R. (2005). Recombination regulation by transcription-induced cohesin dissociation in rDNA repeats. Science 309, 1581–1584. 10.1126/science.1116102.

Kobayashi, T., Heck, D.J., Nomura, M., and Horiuchi, T. (1998). Expansion and contraction of ribosomal DNA repeats in Saccharomyces cerevisiae: requirement of replication fork blocking (Fob1) protein and the role of RNA polymerase I. Genes & Development 12, 3821–3830. 10.1101/gad.12.24.3821.

Kobayashi, T., Hidaka, M., Nishizawa, M., and Horiuchi, T. (1992). Identification of a site required for DNA replication fork blocking activity in the rRNA gene cluster in Saccharomyces cerevisiae. Mol Gen Genet 233, 355–362.

Kobayashi, T., and Horiuchi, T. (1996). A yeast gene product, Fob1 protein, required for both replication fork blocking and recombinational hotspot activities. Genes to cells : devoted to molecular & cellular mechanisms 1, 465–474.

Kobayashi, T., Horiuchi, T., Tongaonkar, P., Vu, L., and Nomura, M. (2004). SIR2 regulates recombination between different rDNA repeats, but not recombination within individual rRNA genes in yeast. Cell 117, 441–453.

Kobayashi, T., and Sasaki, M. (2017). rDNA stability is supported by many “Buffer genes” - Introduction to the Yeast rDNA Stability Database. FEMS Yeast Res. 10.1093/femsyr/fox001.

Kwan, E.X., Wang, X.S., Amemiya, H.M., Brewer, B.J., and Raghuraman, M.K. (2016). rDNA Copy Number Variants Are Frequent Passenger Mutations in Saccharomyces cerevisiae Deletion Collections and de Novo Transformants. G3 (Bethesda) 6, 2829–2838. 10.1534/g3.116.030296.

Laloraya, S., Guacci, V., and Koshland, D. (2000). Chromosomal addresses of the cohesin component Mcd1p. J Cell Biol 151, 1047–1056. 10.1083/jcb.151.5.1047.

Lambert, S., Watson, A., Sheedy, D.M., Martin, B., and Carr, A.M. (2005). Gross chromosomal rearrangements and elevated recombination at an inducible site-specific replication fork barrier. Cell 121, 689–702. 10.1016/j.cell.2005.03.022.

Loberg, M.A., Hurtig, J.E., Graff, A.H., Allan, K.M., Buchan, J.A., Spencer, M.K., Kelly, J.E., Clodfelter, J.E., Morano, K.A., Lowther, W.T., and West, J.D. (2019). Aromatic Residues at the Dimer-Dimer Interface in the Peroxiredoxin Tsa1 Facilitate Decamer Formation and Biological Function. Chem Res Toxicol 32, 474–483. 10.1021/acs.chemrestox.8b00346.

López-Otín, C., Blasco, M.A., Partridge, L., Serrano, M., and Kroemer, G. (2013). The Hallmarks of Aging. Cell 153, 1194–1217. 10.1016/j.cell.2013.05.039.

McHugh, D., and Gil, J. (2018). Senescence and aging: Causes, consequences, and therapeutic avenues. J Cell Biol 217, 65–77. 10.1083/jcb.201708092.

Menzel, J., Malo, M.E., Chan, C., Prusinkiewicz, M., Arnason, T.G., and Harkness, T.A. (2014). The anaphase promoting complex regulates yeast lifespan and rDNA stability by targeting Fob1 for degradation. Genetics 196, 693–709. 10.1534/genetics.113.158949.

Mizuno, K., Miyabe, I., Schalbetter, S.A., Carr, A.M., and Murray, J.M. (2013). Recombination-restarted replication makes inverted chromosome fusions at inverted repeats. Nature 493, 246–249. 10.1038/nature11676.

Molin, M., Yang, J., Hanzen, S., Toledano, M.B., Labarre, J., and Nystrom, T. (2011). Life span extension and H(2)O(2) resistance elicited by caloric restriction require the peroxiredoxin Tsa1 in Saccharomyces cerevisiae. Molecular cell 43, 823–833. 10.1016/j.molcel.2011.07.027.

Mortimer, R.K., and Johnston, J.R. (1959). Life span of individual yeast cells. Nature 183, 1751–1752. 10.1038/1831751a0.

Nystrom, T., Yang, J., and Molin, M. (2012). Peroxiredoxins, gerontogenes linking aging to genome instability and cancer. Genes Dev 26, 2001–2008. 10.1101/gad.200006.112.

Park, S.G., Cha, M.K., Jeong, W., and Kim, I.H. (2000). Distinct physiological functions of thiol peroxidase isoenzymes in Saccharomyces cerevisiae. The Journal of biological chemistry 275, 5723–5732. 10.1074/jbc.275.8.5723.

Perkins, A., Nelson, K.J., Parsonage, D., Poole, L.B., and Karplus, P.A. (2015). Peroxiredoxins: guardians against oxidative stress and modulators of peroxide signaling. Trends Biochem Sci 40, 435–445. 10.1016/j.tibs.2015.05.001.

Perrone, G.G., Tan, S.X., and Dawes, I.W. (2008). Reactive oxygen species and yeast apoptosis. Biochim Biophys Acta 1783, 1354–1368. 10.1016/j.bbamcr.2008.01.023.

Roger, F., Picazo, C., Reiter, W., Libiad, M., Asami, C., Hanzen, S., Gao, C., Lagniel, G., Welkenhuysen, N., Labarre, J., et al. (2020). Peroxiredoxin promotes longevity and H(2)O(2)-resistance in yeast through redox-modulation of protein kinase A. Elife 9. 10.7554/eLife.60346.

Saka, K., Ide, S., Ganley, A.R., and Kobayashi, T. (2013). Cellular senescence in yeast is regulated by rDNA noncoding transcription. Curr Biol 23, 1794–1798. 10.1016/j.cub.2013.07.048.

Saka, K., Takahashi, A., Sasaki, M., and Kobayashi, T. (2016). More than 10% of yeast genes are related to genome stability and influence cellular senescence via rDNA maintenance. Nucleic Acids Res. 10.1093/nar/gkw110.

Sasaki, M., and Kobayashi, T. (2017). Ctf4 Prevents Genome Rearrangements by Suppressing DNA Double-Strand Break Formation and Its End Resection at Arrested Replication Forks. Molecular cell 66, 533–545 e535. 10.1016/j.molcel.2017.04.020.

Sasaki, M., and Kobayashi, T. (2021). Gel Electrophoresis Analysis of rDNA Instability in Saccharomyces cerevisiae. Methods Mol Biol 2153, 403–425. 10.1007/978-1-0716-0644-5_28.

Sasaki, M., and Kobayashi, T. (2023a). Regulatory processes that maintain or alter ribosomal DNA stability during the repair of programmed DNA double-strand breaks. Genes Genet Syst 98, 103–119. 10.1266/ggs.22-00046.

Sasaki, M., and Kobayashi, T. (2023b). Transcription near arrested DNA replication forks triggers ribosomal DNA copy number changes. bioRxiv, 2023.2012.2021.572944. 10.1101/2023.12.21.572944.

Sasaki, M., Lange, J., and Keeney, S. (2010). Genome destabilization by homologous recombination in the germ line. Nat Rev Mol Cell Biol 11, 182–195. 10.1038/nrm2849.

Seo, A.Y., Joseph, A.M., Dutta, D., Hwang, J.C., Aris, J.P., and Leeuwenburgh, C. (2010). New insights into the role of mitochondria in aging: mitochondrial dynamics and more. J Cell Sci 123, 2533–2542. 10.1242/jcs.070490.

Sies, H., and Jones, D.P. (2020). Reactive oxygen species (ROS) as pleiotropic physiological signalling agents. Nat Rev Mol Cell Biol 21, 363–383. 10.1038/s41580-020-0230-3.

Sinclair, D.A., and Guarente, L. (1997). Extrachromosomal rDNA Circles— A Cause of Aging in Yeast. Cell 91, 1033–1042. 10.1016/s0092-8674(00)80493-6.

Sinclair, D.A., Mills, K., and Guarente, L. (1997). Accelerated aging and nucleolar fragmentation in yeast sgs1 mutants. Science 277, 1313–1316. 10.1126/science.277.5330.1313.

Sobotta, M.C., Liou, W., Stocker, S., Talwar, D., Oehler, M., Ruppert, T., Scharf, A.N., and Dick, T.P. (2015). Peroxiredoxin-2 and STAT3 form a redox relay for H2O2 signaling. Nat Chem Biol 11, 64–70. 10.1038/nchembio.1695.

Tachibana, T., Okazaki, S., Murayama, A., Naganuma, A., Nomoto, A., and Kuge, S. (2009). A major peroxiredoxin-induced activation of Yap1 transcription factor is mediated by reduction-sensitive disulfide bonds and reveals a low level of transcriptional activation. The Journal of biological chemistry 284, 4464–4472. 10.1074/jbc.M807583200.

Takeuchi, Y., Horiuchi, T., and Kobayashi, T. (2003). Transcription-dependent recombination and the role of fork collision in yeast rDNA. Genes Dev 17, 1497–1506. 10.1101/gad.1085403.

Thorpe, G.W., Fong, C.S., Alic, N., Higgins, V.J., and Dawes, I.W. (2004). Cells have distinct mechanisms to maintain protection against different reactive oxygen species: oxidative-stress-response genes. Proceedings of the National Academy of Sciences of the United States of America 101, 6564–6569. 10.1073/pnas.0305888101.

Weitao, T., Budd, M., Hoopes, L.L., and Campbell, J.L. (2003). Dna2 helicase/nuclease causes replicative fork stalling and double-strand breaks in the ribosomal DNA of Saccharomyces cerevisiae. The Journal of biological chemistry 278, 22513–22522. 10.1074/jbc.M301610200.

Wong, C.M., Siu, K.L., and Jin, D.Y. (2004). Peroxiredoxin-null yeast cells are hypersensitive to oxidative stress and are genomically unstable. The Journal of biological chemistry 279, 23207–23213. 10.1074/jbc.M402095200.

Wong, C.M., Zhou, Y., Ng, R.W., Kung Hf, H.F., and Jin, D.Y. (2002). Cooperation of yeast peroxiredoxins Tsa1p and Tsa2p in the cellular defense against oxidative and nitrosative stress. The Journal of biological chemistry 277, 5385–5394. 10.1074/jbc.M106846200.

Wood, Z.A., Schroder, E., Robin Harris, J., and Poole, L.B. (2003). Structure, mechanism and regulation of peroxiredoxins. Trends Biochem Sci 28, 32–40. 10.1016/s0968-0004(02)00003-8.

Yokoyama, M., Sasaki, M., and Kobayashi, T. (2023). Spt4 promotes cellular senescence by activating non-coding RNA transcription in ribosomal RNA gene clusters. Cell Rep 42, 111944. 10.1016/j.celrep.2022.111944.

