## Supplemental Table 1 for "The peroxiredoxin Tsa1 extends the lifespan of budding yeast by maintaining the stability of the ribosomal RNA gene cluster"

**Supplementary Table 1.** ***S. cerevisiae*** **strains used in this study**

| Name | Genotype |
|---|---|
| JOY100 | |
| JOY102 | *sch9Δ::kanMX* |
| JOY103 | *vps73Δ::kanMX* |
| JOY104 | *sod1Δ::kanMX* |
| JOY105 | *rad23Δ::kanMX* |
| JOY106 | *atp5Δ::kanMX* |
| JOY108 | *ypr124wΔ::kanMX* |
| JOY109 | *ymr038cΔ::kanMX* |
| JOY111 | *slf1Δ::kanMX* |
| JOY112 | *gcr6Δ::kanMX* |
| JOY113 | *hsv2Δ::kanMX* |
| JOY114 | *ssq1Δ::kanMX* |
| JOY115 | *pdx3Δ::kanMX* |
| JOY118 | *abz1Δ::kanMX* |
| JOY119 | *vma22Δ::kanMX* |
| JOY120 | *pth1Δ::kanMX* |
| JOY123 | *hsp78Δ::kanMX* |
| JOY124 | *jen1Δ::kanMX* |
| JOY125 | *rtc6Δ::kanMX* |
| JOY126 | *mid1Δ::kanMX* |
| JOY127 | *tda1Δ::kanMX* |
| JOY128 | *adh2Δ::kanMX* |
| JOY131 | *sec28Δ::kanMX* |
| JOY133 | *aim19Δ::kanMX* |
| JOY134 | *oar1Δ::kanMX* |
| JOY135 | *idp2Δ::kanMX* |
| JOY136 | *bts1Δ::kanMX* |
| JOY138 | *arg4Δ::kanMX* |
| JOY139 | *lys9Δ::kanMX* |
| JOY140 | *met22Δ::kanMX* |
| JOY141 | *trp1Δ::kanMX* |
| JOY142 | *mal31Δ::kanMX* |
| JOY143 | *aro3Δ::kanMX* |
| JOY144 | *ape3Δ::kanMX* |
| JOY145 | *zrc1Δ::kanMX* |
| JOY146 | *hrb1Δ::kanMX* |
| JOY147 | *lst4Δ::kanMX* |
| JOY148 | *pxa1Δ::kanMX* |
| JOY149 | *ubp2Δ::kanMX* |
| JOY152 | *aim10Δ::kanMX* |
| JOY153 | *coq6Δ::kanMX* |
| JOY154 | *ccw14Δ::kanMX* |
| JOY155 | *arg2Δ::kanMX* |
| JOY156 | *ypr099cΔ::kanMX* |
| JOY157 | *mrm2Δ::kanMX* |
| JOY158 | *mtg2Δ::kanMX* |
| JOY159 | *trm10Δ::kanMX* |

| | |
|---|---|
| JOY160 | *ydl086wΔ::kanMX* |
| JOY161 | *ynl144cΔ::kanMX* |
| JOY162 | *voa1Δ::kanMX* |
| JOY163 | *pir5Δ::kanMX* |
| JOY164 | *ycl021w-aΔ::kanMX* |
| JOY165 | *ser3Δ::kanMX* |
| JOY166 | *lys7Δ::kanMX* |
| JOY167 | *skn7Δ::kanMX* |
| JOY168 | *yap1Δ::kanMX* |
| JOY169 | *ahp1Δ::kanMX* |
| JOY170 | *prx1Δ::kanMX* |
| JOY171 | *dot5Δ::kanMX* |
| JOY172 | *sod2Δ::kanMX* |
| JOY173 | *trx1Δ::kanMX* |
| JOY174 | *trx2Δ::kanMX* |
| JOY175 | *trx3Δ::kanMX* |
| JOY176 | *trr2Δ::kanMX* |
| JOY177 | *gpx1Δ::kanMX* |
| JOY178 | *gpx2Δ::kanMX* |
| JOY179 | *hyr1Δ::kanMX* |
| JOY180 | *gsh1Δ::kanMX* |
| JOY181 | *gsh2Δ::kanMX* |
| JOY182 | *ccp1Δ::kanMX* |
| JOY183 | *ctt1Δ::kanMX* |
| JOY184 | *cta1Δ::kanMX* |
| JOY186 | *tkl1Δ::kanMX* |
| JOY187 | *rpe1Δ::kanMX* |
| JOY188 | *elg1Δ::kanMX* |
| JOY211 | *rad27Δ::kanMX* |
| JOY256 | *tsa2Δ::kanMX* |
| JOY268 | *tsa1Δ::kanMX/TSA1, fob1::LEU2/FOB1* |
| JOY263 | *tsa1Δ::hphMX/TSA1, tsa2Δ::kanMX/TSA2* |
| JOY275 | YEp112 |
| JOY276 | YEplac112-TSA1 |
| JOY277 | *tsa1Δ::hphMX*, YCplac22 |
| JOY278 | *tsa1Δ::hphMX*, YCplac22-TSA1 |
| JOY279 | *tsa1Δ::hphMX,* YEplac112 |
| JOY280 | *tsa1Δ::hphMX,* YEplac112-TSA1 |
| JOY283 | *sir2Δ::kanMX*, YEplac112 |
| JOY284 | *sir2Δ::kanMX*, YEplac112-TSA1 |
| JOY285 | *tsa1Δ::hphMX*, YCplac22-*tsa1*$^{C48S}$ |
| JOY286 | *tsa1Δ::hphMX*, YCplac22-*tsa1*$^{C171S}$ |
| JOY287 | *tsa1Δ::hphMX,* YCplac22-*tsa1*$^{F44A}$ |
| JOY288 | *tsa1Δ::hphMX,* YCplac22-*tsa*$^{Y78A}$ |
| JOY290 | *tsa1Δ::hphMX*, YCplac22-*tsa1*$^{C48S,\ C171S}$ |
| JOY294 | *tsa1Δ::hphMX/TSA1, srx1Δ::kanMX/SRX1* |
| JOY295 | *tsa1Δ::hphMX,* YCplac22-*tsa1*$^{F44A,\ Y78A}$ |
| JOY301 | *sir2Δ::kanMX* |
| JOY302 | *fob1::LEU2* |

| | |
|---|---|
| JOY401 | *tsa1Δ::hphMX* |
| MSY1654 | *sir2Δ::hphMX, fob1::LEU2* |

All strains are derivatives of W303, which is *ade2-1, ura3-1, his3-11, 15, trp1-1, leu2-3, 112, can1-100*.
